# Genotype and farm effects on yield and morphology reveal potential for breeding and site selection for sugar kelp

**DOI:** 10.64898/2026.05.10.722392

**Authors:** Sanne Put, Andries Temme, Jessica Schiller, Brigit Reus, Gabriel Montecinos Arismendi, Tijs Ketelaar, Luisa M. Trindade

## Abstract

Seaweed cultivation has recently gained increased attention in North-West Europe as a sustainable source of biomass for biobased products. However, yields need to increase to make the seaweed sector economically viable. To achieve this, higher yielding varieties can be bred but this requires variation for yield and yield-related traits among genotypes. To reliably select high-yielding genotypes, an understanding is required of how both within-farm and between-farm environmental differences affect phenotypes and how to identify simple and reliable proxies for yield. In this study we evaluated growth of nine *Saccharina latissima* genotypes on two farms, 12 km apart, within the same season. We observed a threefold difference in yield among genotypes, demonstrating the potential for improvement through selection and breeding. Blade thickness and blade size-related traits were strongly correlated with yield, highlighting their potential to serve as rapid and non-destructive proxies for yield, thereby accelerating selection. Furthermore, we demonstrated the importance of adequate replication in farm trials to improve genotype performance estimation by correcting for within-farm spatial variation. Moreover, phenotypic variation was most explained by the genotype and environment, highlighting the importance of both genotype and site selection. Although genotype by environment interactions (GxE) were significant, its contributions were small, indicating stable genotype ranking across farms. Overall, these results are promising for breeding improved *S. latissima* as it indicates that genotype performance is consistent across close by locations and that local *S. latissima* populations harbour substantial phenotypic variation that can be used to breed for increased yield.

**Highlights:**

- Local genetic resources harbour substantial variation in yield and morphology for breeding.
- Minor GxE allows for breeding across farms.
- Blade thickness and blade size related traits are good predictors of yield.
- Correction for on-farm spatial variation improves genotype performance estimation.

## 1. Introduction

Currently, seas in North-West Europe are underutilized for sustainable biomass production. Seaweed production could aid in transitioning towards a biobased economy with locally and sustainably farmed biomass (EC, 2022). For North-West Europe specifically, *Saccharina latissima* is a species with high potential. First, *S. latissima* is a native species along the European Atlantic coastline, which is important in seaweed cultivation to prevent the cultivation of non-native species. Second, it is one of the highest-yielding seaweed species native to North-West Europe (Fernand et al., 2017; Kraan, 2013; Van Der Molen et al., 2018). Nevertheless, further increases in yields are required to make the seaweed farming sector economically viable (van den Burg et al., 2016).

Breeding for increased yield has been successful in closely related seaweed species. For example, *Saccharina japonica* has achieved large yield increases since the initiation of breeding programs in the 1950s (Hu et al., 2021; X. Li et al., 2007; Wang et al., 2020; Zhang et al., 2007). Yet, only limited breeding efforts have been made for increased yield *S. latissima*, suggesting considerable potential for genetic improvement (Huang et al., 2022). In selection and breeding, a major advantage of *S. latissima* is its biphasic life cycle with distinct alternating haploid gametophyte and diploid sporophyte generations (Kanda, 1936). The sporophytes are large and represent the harvestable life stage, whereas the gametophytes are much smaller. Haploid gametophytes can be clonally propagated, which enables crossing clonal gametophytes to obtain genetically identical sporophytes within a single generation (D. Li et al., 1999; Robinson et al., 2013). This improves genotype performance evaluation and reduces the many generations of selfing, often needed in plant breeding to obtain uniform lines.

Despite this opportunity, breeding for yield is complex. Yield is influenced by both the genotype, the environment, and a combination of the two. In addition, yield depends on other traits, such as density, blade size, and blade thickness (Bråtelund et al., 2026; Cohen et al., 2025; Gerard et al., 1987; Gonzalez et al., 2025; Huang et al., 2023). Identifying and verifying easy-to-measure proxy traits is necessary to overcome barriers of limited farm access during the season, high costs of sea-based operations and the fast deterioration of sporophytes after harvest. As such, gaining an understanding of the trait relationships can simplify and facilitate breeding (Robinson et al., 2013).

Several studies have demonstrated the effect of different environments on yield and morphology of a single genotype of *S. latissima*. For example, stronger currents result in longer and heavier blades with a higher length-to-width ratio (Buck & Buchholz, 2005; Gerard, 1987; Peteiro & Freire, 2013). Thicker, longer and narrower blades experience less drag, making them more robust and less prone to dislodgement at exposed sites. Additionally, increased light intensities and a higher temperature and nutrient availability result in longer blades and a faster growth (Bruhn et al., 2016; Chopin et al., 2004; Diehl & Bischof, 2021; Iñiguez et al., 2016; Sanderson et al., 2008; Vilmin & Van Duren, 2021). Light intensity can vary due to cultivation depth and water turbidity, the latter affected by seabed depth and softness. Even though *S. latissima* is known to be plastic and thus changes morphology across environments, less is known about trait correlations for different genotypes across environments. A better understanding of trait correlations across environments would enable the identification of traits that are both predictive of yield and quick and easy to evaluate.

Although genotype differences in morphology and yield have been studied within a single environment, multi-environmental comparisons are still limited (Bråtelund et al., 2026; Cohen et al., 2025; Huang et al., 2023). Because of environmental protection measures *S. latissima* grown on farms must originate from local populations. Thus, differential ranking of genotypes across farms requires location specific breeding programs, which are costly and time-intensive (Robinson et al., 2013). Understanding the extent of genotype by environment interaction (G×E) for different traits is therefore essential. However, genotype comparisons across farms have been limited. Recently, Gonzalez et al. (2025) evaluated six genotypes across three farms and observed differential performance of genotypes across these farms for yield traits but not for blade morphology. The farms in their study were relatively far apart, with approximately 300 km separating the most distant farms. The question remains whether genotypes also rank differently between geographically closer farms, where environmental differences may be smaller.

Besides variation in environmental conditions between farms, conditions can also vary within farms. This spatial heterogeneity, potentially reflecting local differences in water flow, nutrient availability and light conditions, can influence yield. For instance, Bråtelund et al. (2025) observed a tendency for an increase in yield along the sequence of cultivation ropes, which suggests an effect of positioning within the farm. This offers the opportunity to improve genotype performance estimation by modelling environmental variation (Araus & Cairns, 2014; Ball et al., 1993). In contrast to agricultural breeding trials, within-farm environmental variability is still poorly understood in *S. latissima* breeding trials.

In this study we aim to understand the extent of variability in yield of *S. latissima* across different growth environments, different genotypes and their combination. Moreover, we aim to determine which morphological traits are important in selection for yield and how these traits are affected by the environment. We addressed the following questions: (1) How do genotype, environment, and their interaction affect yield, morphology and yield-related traits on farms that are relatively close to each other? (2) Which morphological traits are correlated with yield? (3) Can accounting for within-farm spatial trends improve genotype performance estimation? To do so, we conducted two trials across relatively proximate farms during the same growing season.

## 2. Materials and Methods

### 2.1. Seaweed material

To generate seeding material, clonal gametophyte cultures were obtained from individual sporulating sporophytes collected at three different locations in the Vestland area, Norway (Table S1). Spore extraction and gametophyte clonal propagation were performed by Hortimare using standard procedures, including sterile conditions, gametophyte isolation, temperature, and light control conditions (Ebbing et al., 2021). The gametophyte cultures were maintained at 12°C and 5 µmol photons m−2 s −1 red light and a 16-hour photoperiod (Theodorou et al., 2021). Twelve different crosses were made with a clonal female and male gametophyte culture that originate from the same parental sporophytes. Before making crosses, female and male gametophytes were separately fragmented with Potter–Elvehjem tissue grinders and combined at a 1:1 ratio. Gametogenesis was induced by cultivating the combined cultures at 10°C and 50 µmol photons m−2 s −1 white light and a 14-hour photoperiod (Theodorou et al., 2021). After 15/16 days, sporophytes were sown on polyester twine with an initial density of 1600/2000 sporophytes per meter. Spools were transported in temperature control condition by Hortimare. Sporophytes attached to twine reached 3-5 mm of length at moment of deployment at the farm.

### 2.2. Farm description, deployment and harvest

Polyester twine with seedlings was wound around cultivation lines and deployed on October 26^th^, 2023, and October 27^th^, 2023, for the farms Lerøy Ocean Harvest (hereafter: Leroy/Lerøy) (60.148278 N, 5.235500 E) and Austevoll Sea Organic AS (hereafter: Austevoll) (60.040639, 5.192056) respectively. This corresponds to 45 days and 46 days after combining female and male gametophytes. Out of the 12 crosses, 11 crosses were grown on both farms. Section numbers follow the ropes and start on the bottom left (X and Y value of 0). Due to a limited availability of twine, we did not deploy cross number 10 at Austevoll and on both farms a number of replicates between 5 and 8 was used for all crosses (Table S1). Each seeded rope segment (replicate) was 1 meter long with 0.5-meter unseeded rope between the segments. Genotype replicates were randomly distributed over the farms among other crosses, not part of this study, that were also grown on the farms (Figure 1). Harvesting was done on the 15^th^ of April 2025 for Lerøy and on the 22^nd^ of April 2025 for Austevoll. During the harvest, the cultivation lines were lifted out of the water and pictures were taken of the whole 1-meter section. Afterwards, sporophytes were collected from a 20 cm section, stored in mesh bags and transported to land. The bags were either kept in large containers with running seawater or in the sea until phenotyping, which took place within two days after harvest.

**Figure 1.**
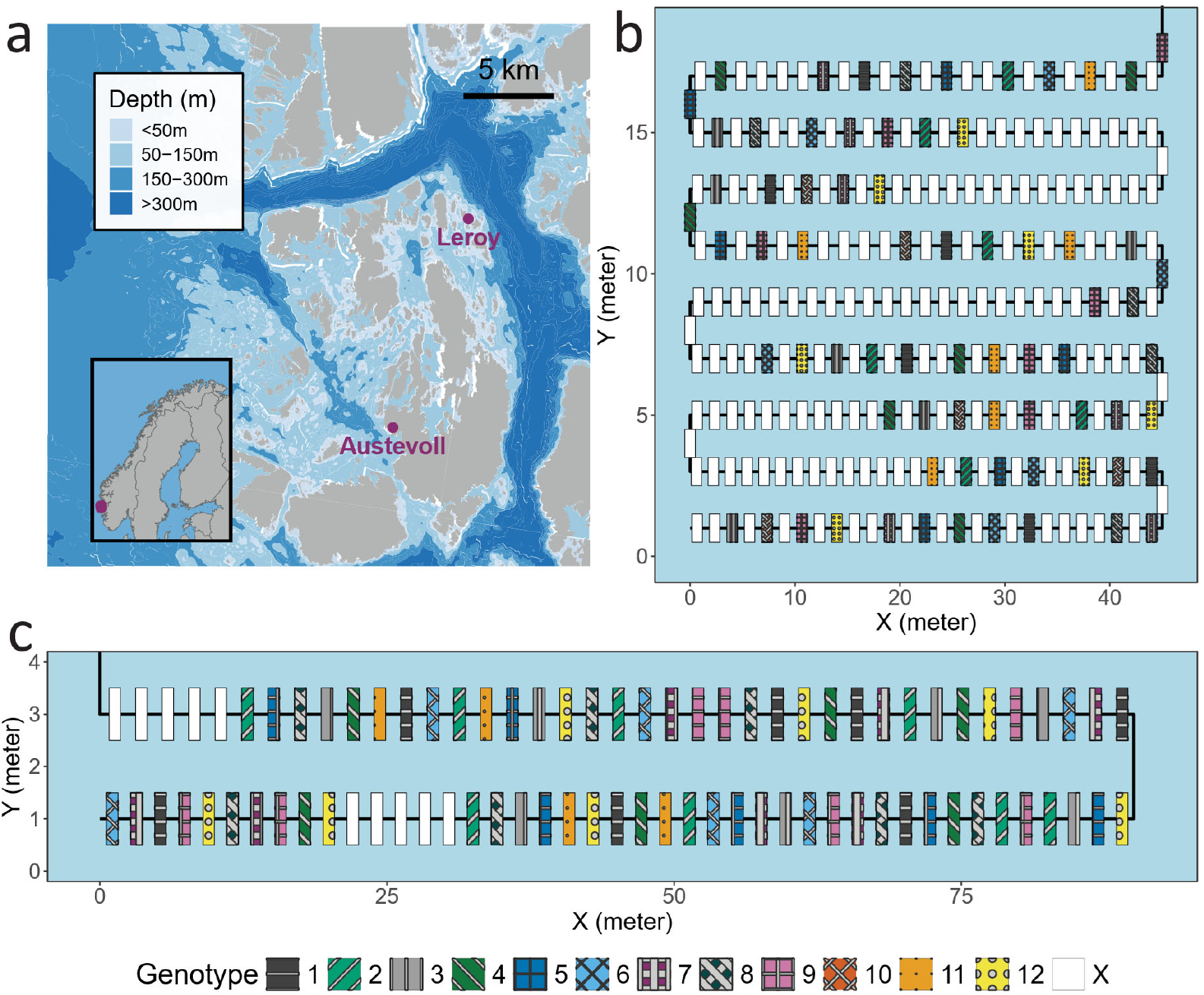
Farm location and design. (a) Map of farm locations. Farms are indicated in red and sea depths are indicated by shades of blue (Kartverket, n.d.). (b) Farm design of Lerøy. (c) Farm design of Austevoll. For both farm designs more sections follow the rope outside the figure that were not included in this experiment.

### 2.3. Phenotyping

For each harvested section, total sporophyte fresh weight and the number of sporophytes (density_20cm_) longer than 15 cm were recorded. For all sections at Lerøy and the first three sections at Austevoll, 8 sporophytes were randomly selected for morphological blade measurements. Blades were laid flat and imaged with a 25cm ruler included as scale reference. Additionally, one blade was randomly selected for measurement of thickness and individual blade fresh and dry weight. Thickness measurements were performed using an electronic thickness gauge at the bottom and the middle along the blade length, measured at the centre of the blade width. Imaged sporophyte blades were first air- and sun-dried, then subsequently dried at 40°C for 72 hours prior to dry weight determination. Blade morphological traits were determined using ImageJ (Schindelin et al., 2012). For Austevoll, the camera was not positioned exactly horizontally and centred above the sporophytes. Therefore, the corners of the tables were used to change the perspective of the image using the Perspective Wrap function in Adobe Photoshop 2024 (Adobe Inc., San Jose, CA, USA). This was not needed for Lerøy because the camera was positioned horizontally and centred above the sporophytes. The morphological characteristics measured were blade area, perimeter, circularity, aspect ratio (AR), roundness, solidity, length and maximum width (Figure 2). The circularity was calculated by (4π × area)/(perimeter)^2^, roundness by (4 × area)/(4π × (length^2^), solidity by area/convex area and the aspect ratio by length/width. Circularity, roundness and solidity range from 0 to 1.

**Figure 2.**
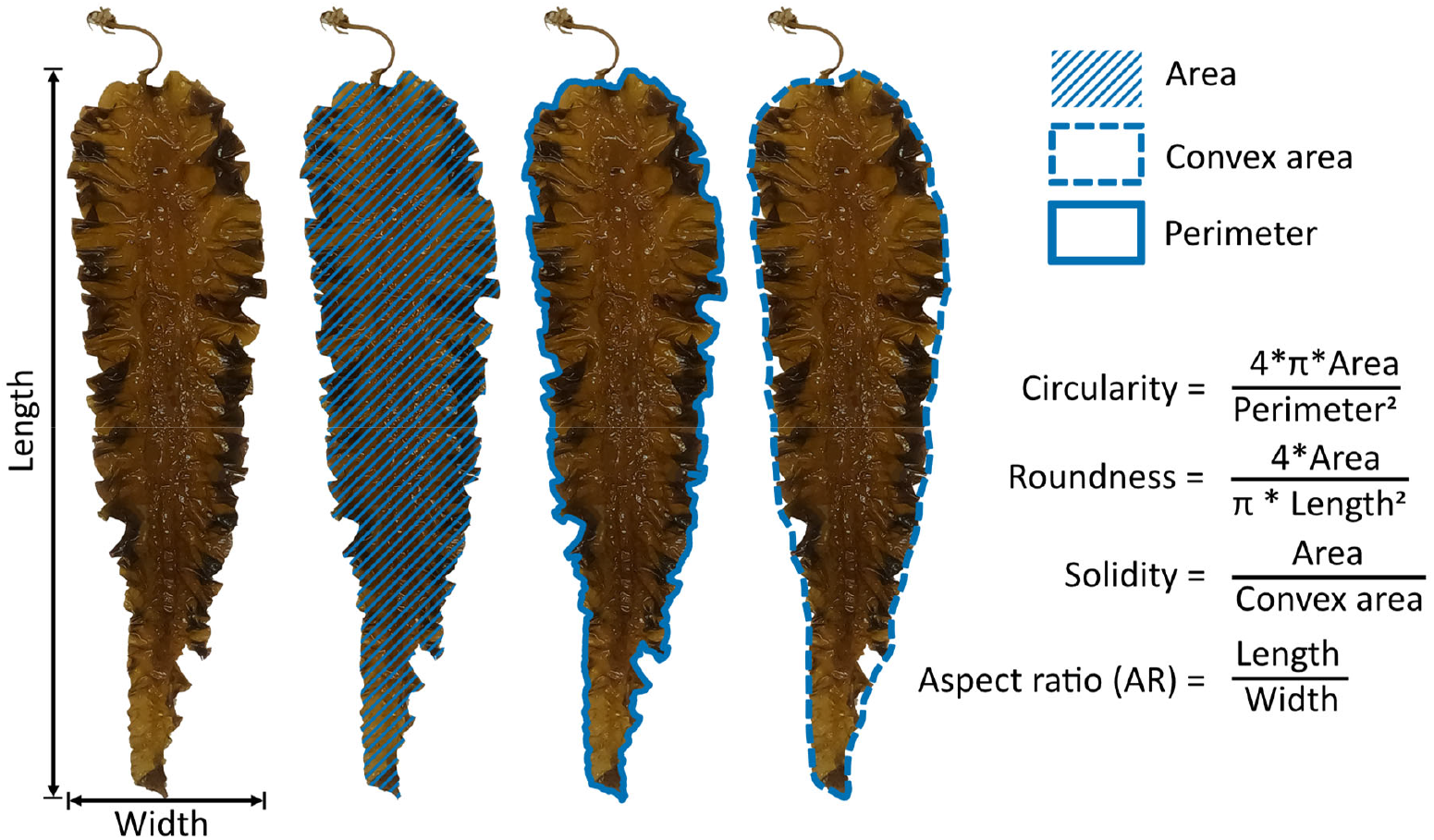
Blade shape traits. The convex area is the area covered by the convex hull. Circularity is calculated by (4π × area)/(perimeter)2, roundness by (4 × area)/(4π × (length2), solidity by area/convex area and the aspect ratio by length/width.

### 2.4. Statistical analysis

Results were analysed using R (v4.4.1, R Core Team, 2024) in RStudio (RStudio Team, 2024). All plots were made with ggplot2 (version 3.5.2). Traits bounded between 0 and 1 (circularity, roundness and solidity) were logit-transformed prior to statistical analysis to meet assumptions (Warton & Hui, 2011).

Spatial correction was performed based on 2D P-splines for Lerøy with SpATS (Rodríguez-Álvarez et al., 2018). For Austevoll, LMMSolver (Boer, 2023) was used based on 1D P-splines due to absence of a 2D spatial trend (only two rows), with the rows (Y) as fixed effect. To fit the splines, the numbers of segments used were chosen based on approximately 1 knot per plot in each spatial dimension. This resulted in 20 segments in the X-direction and 8 segments in the Y-direction for Lerøy and 40 segments in the X-direction for Austevoll. Penalization prevented overfitting, even with high numbers of segments. Fitted values were obtained from the models including genotype as fixed effect. Farm specific generalized heritabilities and variance components were extracted using genotype as a random effect (Oakey et al., 2006). For combined analysis across farms, generalized heritability was estimated with genotype and GxE effects modelled as random. GxE analysis was performed using a linear model including genotype, farm and their interaction, and the effects were considered significant for p < 0.05. Sidak-adjusted pairwise comparisons of estimated marginal means were used to determine differences among genotypes within each farm, while Tukey-adjusted comparisons were used to test for differences among farms within each genotype (Šidák, 1967; Tukey, 1949). A significance threshold of p < 0.05 was used for all tests.

To study trait relationships, Spearman rank correlations for the farms separately were calculated using genotype means (Spearman, 1904). Trait correlations for the farms combined were calculated with genotype means corrected for GxE to prevent Simpson’s paradox (Simpson, 1951). Principal Component Analysis (PCA) was performed using the prcomp package on scaled and centred trait values, including only the plots with complete observations.

## 3. Results

### 3.1. Genotypes differ substantially for yield and morphology

To determine the potential for breeding of *S. latissima* the phenotypic diversity between genotypes was quantified. We observed a large diversity between genotypes, both for the whole 1m section (Figure 3A), as well as for the morphology of randomly selected blades (Figure 3B). These examples illustrate the diversity observed at Lerøy (see full variation in Figure S1). This showcases the large phenotypic variation for both yield and morphology between genotypes that can be exploited. Due to exceptionally poor performance, expected to be caused by a disease (wrinkly phenotype), we decided to exclude cross 2 and cross 8 from subsequent analysis (Figure S2).

**Figure 3.**
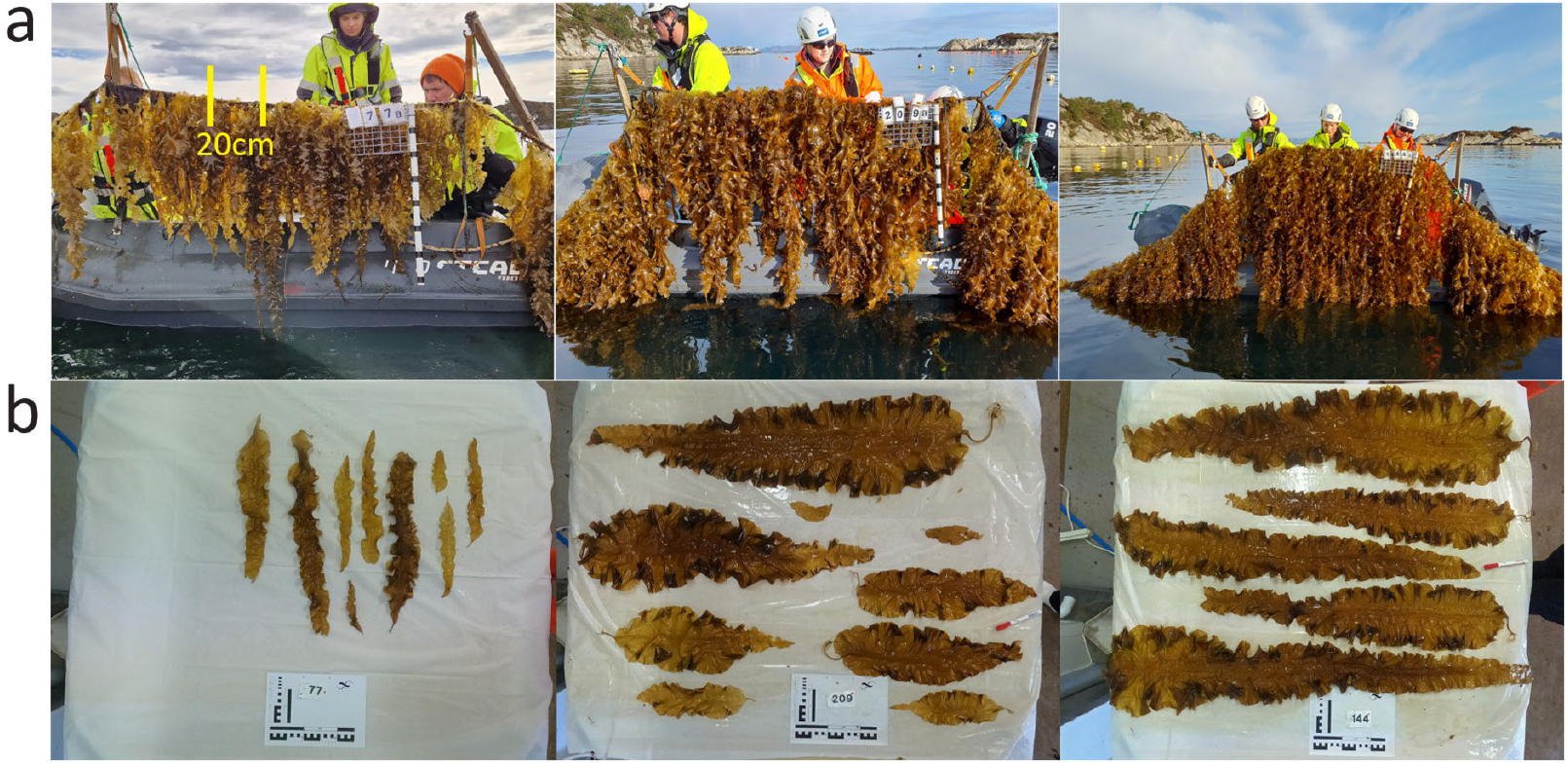
Representation of phenotypic variation between plots. Example of sampling section indicated in green. (a) Plots at Lerøy, from left to right genotype 9, 6 and 11. (b) Morphology of randomly selected blades from the same section as the images in A at Lerøy. From left to right genotype 9, 6 and 11.

### 3.2. Genotype performance for yield is affected by the farm

To assess the effect of the environment and genotype on performance, we evaluated 10 genotypes on two farms. A large phenotypic diversity among genotypes was observed at both farms (Figure 4A). The wet-weight of the sampled 20 cm (wet-weight_20cm_) ranged from 180g (genotype 9) to 1029g (genotype 11), representing a three-fold difference among genotype means. Seven out of ten genotypes significantly differed for yield between the farms. Most genotypes obtained a higher wet-weight_20cm_ in Lerøy compared to Austevoll, except for genotypes 6 and 9. Consequently, a genotype by environment interaction (GxE) was detected for wet-weight_20cm_, although the interaction was primarily driven by a few genotypes behaving in a different manner across environments. Both genotype (G) and environment (E) effects were significant (Figure 4B). Within each environment, genotypes were separated into statistically distinct groups based on a post-hoc comparison (α < 0.05). Clearer genotype separation was obtained at Lerøy than at Austevoll due to lower within-genotype variation at Lerøy.

**Figure 4.**
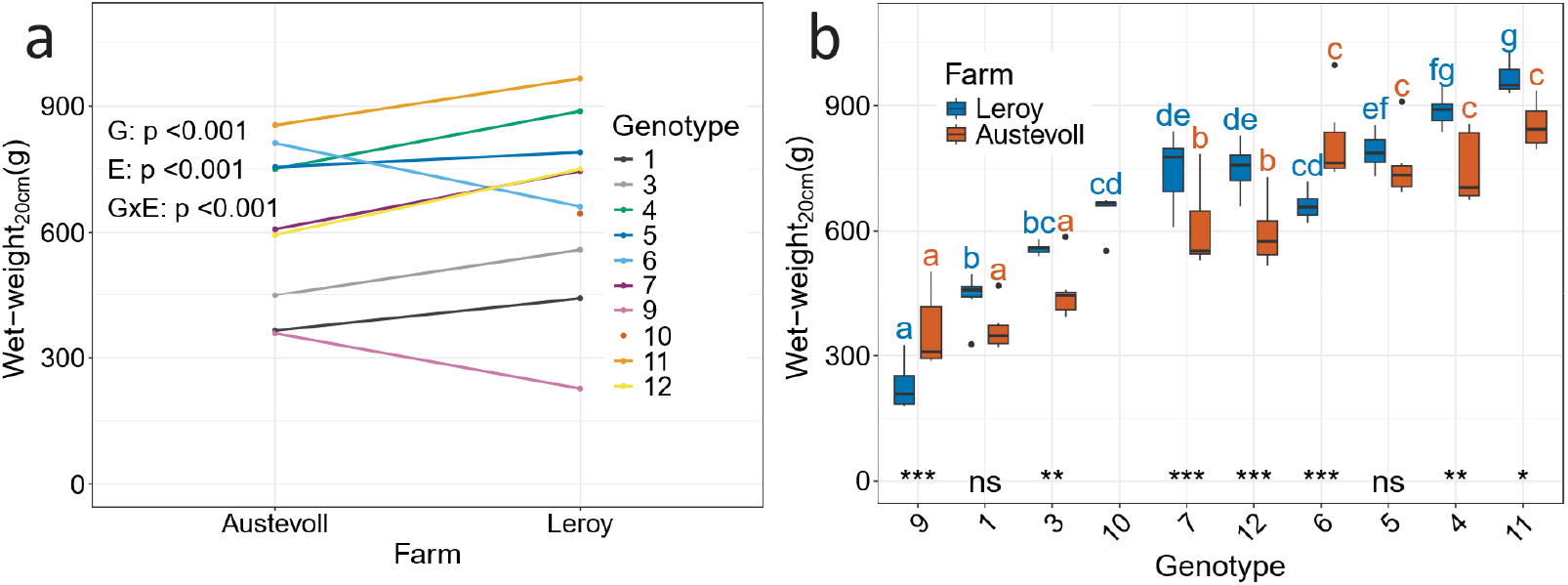
Genotype and environment effect, and their interaction (GxE) for wet-weight_20cm_ for 10 genotypes, of which 9 were grown in both farms. (a) GxE interaction plot with significance derived from a linear mixed model testing the effects of genotype (G), environment (E) and their interaction (GxE). (b) Boxplot with genotype groups within each environment based on Sidak-adjusted post-hoc test (α=0.05). Asterisk at the bottom indicates significance of the difference in performance for a genotype between the two farms (ns, non-significant; *p<0.05; **p<0.01; ***p<0.001).

### 3.3. Stable performance among farms for genotypes

To further study the phenotypic variation among genotypes and farms we explored different morphological traits. On both farms, each trait exhibited substantial variation, with overlapping ranges between the farms. On average, wet-weight_20cm_, blade area and other blade size-related traits were higher for Lerøy, whereas density was higher in Austevoll. Most traits followed similar patterns to yield, with significant GxE, G and E effects for nearly all traits (Table 1, Figure S3). The GxE was again primarily driven by a few genotypes, which is also visible in the variance component partitioning (Figure S4). Most of the variance was attributed to the genotype and environment components, with particularly high genotype variance for wet-weight_20cm_, thickness_bottom_, dry matter content and blade shape traits. The proportion of variance attributed to GxE ranged from 0.05 to 0.40 for the different traits. Overall, trait variation was mainly influenced by the genotype and the environment, rather than their interaction.

**Table 1.** Trait values for both farms. Trait averages per farm, the range (in parentheses) and the generalised heritability for the farms combined. Significance for genotype (G), Farm (E) and their interaction (GxE) is indicated with asterisks (ns, non-significant; *p<0.05; **p<0.01; ***p<0.001).

| Trait | Lerøy |  | Austevoll |  | Heritability | G | E | GxE |
| --- | --- | --- | --- | --- | --- | --- | --- | --- |
|  | Average | Range | Average | Range |  |  |  |  |
| Wet-weight<sub>20cm</sub> (g) | 661.75 | (180.19 - 1028.55) | 601.27 | (286.87 - 996.38) | 0.90 | *** | *** | *** |
| Density<sub>20cm</sub> | 23.72 | (12.58 - 47.72) | 33.65 | (16.19 - 55.21) | 0.84 | *** | *** | *** |
| Thickness<sub>bottom</sub> (mm) | 0.7 | (0.44 - 1.11) | 0.74 | (0.38 - 1.16) | 0.66 | *** |  | *** |
| Thickness<sub>middle</sub> (mm) | 0.38 | (0.26 - 0.54) | 0.3 | (0.22 - 0.41) | 0.41 | *** | *** | *** |
| Dry matter content (%) | 0.08 | (0.06 - 0.11) | 0.08 | (0.06 - 0.09) | 0.82 | *** | ** |  |
| Area (cm<sup>2</sup>) | 647.49 | (148.52 - 1072.57) | 443.01 | (193.23 - 892.97) | 0.67 | *** | *** | *** |
| Length (cm) | 65.93 | (44.04 - 100.35) | 57.09 | (35.65 - 81.93) | 0.67 | *** | *** | *** |
| Width (cm) | 13.27 | (5.82 - 18.7) | 10.12 | (5.96 - 15.22) | 0.86 | *** | *** | *** |
| Perimeter (cm) | 195.69 | (111.57 - 302.57) | 162.46 | (99.51 - 258.41) | 0.53 | *** | *** | *** |
| Wet-weight per area (g/cm<sup>2</sup>) | 0.07 | (0.05 - 0.09) | 0.05 | (0.04 - 0.07) | 0.79 | *** | *** | *** |
| Circularity | 0.2 | (0.13 - 0.25) | 0.2 | (0.14 - 0.26) | 0.72 | *** |  | *** |
| Aspect Ratio | 4.65 | (2.91 - 7.51) | 5.3 | (3.21 - 8.16) | 0.97 | *** | *** | *** |
| Roundness | 0.25 | (0.14 - 0.38) | 0.21 | (0.12 - 0.32) | 0.97 | *** | *** | *** |
| Solidity | 0.82 | (0.77 - 0.86) | 0.84 | (0.77 - 0.87) | 0.82 | *** | *** | *** |

To estimate the genetic contribution to trait variation across genotypes and farms, we estimated the genetic heritability of the traits. Across farms, generalized heritability was moderate to high, ranging from 0.41 to 0.97 (Table 1). The generalized heritability was higher for thickness_bottom_ than for thickness_middle_. When analysing the farms separately, generalised heritability at Austevoll was low for the traits thickness, dry matter content, area, length and circularity (Table S2). The low generalised heritability corresponds to the smaller proportion of genetic variance relative to the spatial and residual variance at Austevoll (Figure 5). For these traits, three replicates were measured compared to six replicates at Lerøy. The higher number of replicates at Lerøy improved estimation of and correction for spatial trends, reducing variation among replicates and increasing generalised heritability.

**Figure 5.**
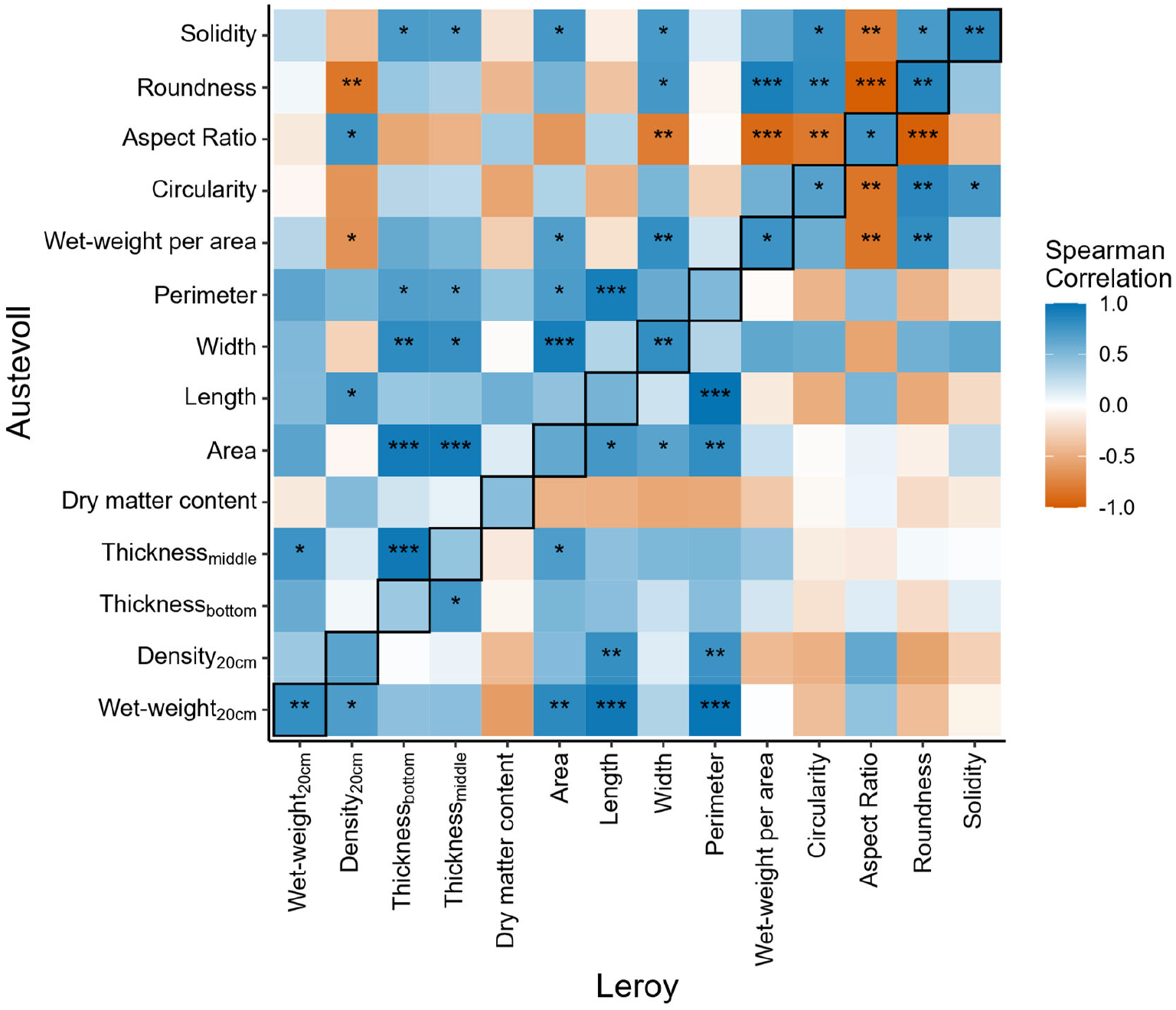
Correlation matrix for traits within the two farms and between the farms. On the diagonal, the correlation for all traits between the two farms is shown. The upper triangle shows the correlation between traits for Austevoll, and the lower triangle shows the correlation between traits for Lerøy. Spearman correlation is indicated by colour ranging from -1 (red) to 1 (blue). Significance is indicated with asterisks (*p<0.05; **p<0.01; ***p<0.001).

### 3.4. Blade thickness, density and blade area-related traits correlate positively with yield

Relationships among traits and their consistency between farms were assessed using correlation analysis. All traits positively correlated between farms, with significant positive correlations for wet-weight_20cm_, length, width, wet-weight per area, circularity, aspect ratio, roundness and solidity (Figure 5). Wet-weight_20cm_ positively correlated on one or both farms with density_20cm_, thickness_middle_, area, length and perimeter. Dry matter content was negative, but non-significantly, correlated with wet-weight_20cm_. The shape traits (circularity, aspect ratio, roundness and solidity) were not significantly correlated with wet-weight_20cm,_ and the magnitude of correlation was low. In contrast, some shape traits did correlate with density_20cm_. For example, density_20cm_ increased with a higher aspect ratio, indicating that a higher density results in longer and narrower sporophytes. Thickness measurements were slightly but non-significantly correlated with wet-weight per area. Furthermore, thickness was not correlated with dry matter content, thus thicker blades likely result in higher dry weight. Overall, trait correlations were consistent between farms, which was also shown when combining the data from both farms (Figure S5).

### 3.5. Moderate multivariate separation of genotypes across farms

To investigate trait correlations among genotypes and farms in a multivariate manner we performed a principal component analysis (PCA). The first two major axes already explained 70% of the total trait variation (Figure 6). The yield trait, wet-weigth_20cm_, was predominantly defined by the second major axis (PC2) and was highly related to blade perimeter and length. In contrast, wet-weight_20cm_ showed little association with the blade shape traits circularity, solidity, and roundness, as well as density_20cm_. The genotypes moderately separated based on the first two principal components (Figure 6A). Genotype 9 clearly separated, mainly driven by PC1. Moreover, genotypes 11 and 12 separated from the rest of the genotypes, predominantly influenced by the density_20cm_ and the aspect ratio. Genotypes 1, 3, 7, and 10 grouped together and were characterised by the blade traits circularity, solidity, and roundness. The last group was composed of genotypes 4, 5, and 6, and was associated with thickness, area, width, length, perimeter and the wet-weight_20cm_. The two farms did not cluster in distinct groups, indicating that they share overall trait variation (Figure 6B). Clustering by genotype rather than environment indicates that genotypes retain their multivariate trait profiles across environments, making these trait profiles useful for selection.

**Figure 6.**
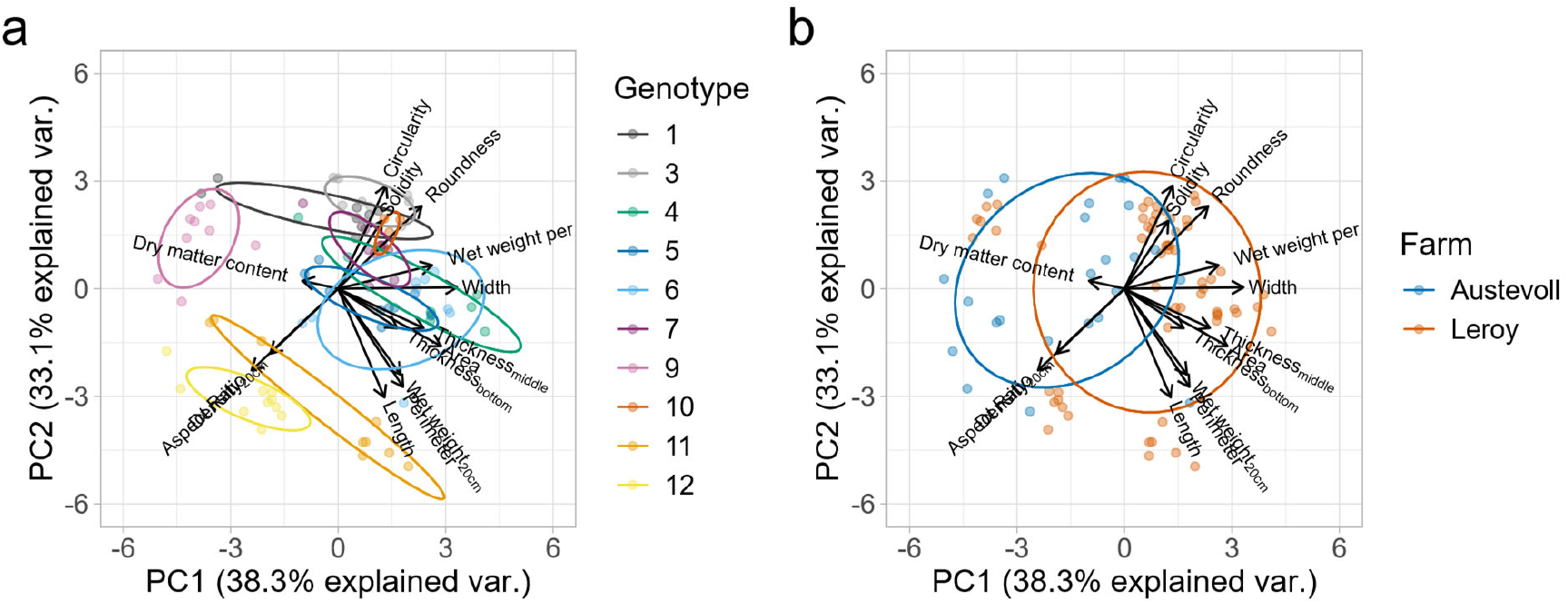
Principal component analysis (PCA) of plot and blade traits. (a) Grouping based on genotype. (b) Grouping based on farm.

### 3.6. Effect of spatial correction depends on the farm and the trait

To determine if within-farm spatial correction improves the estimation of genotype performance, we investigated how the total phenotypic variance was partitioned among the genetic, spatial, and residual components. When a substantial proportion of variance is captured for the spatial trend, the model can capture and correct for the spatial heterogeneity, thereby increasing the precision of the estimated genetic effect. We observed that variance partitioning is different between farms and traits (Figure 7). Wet-weight_20cm_ was only slightly influenced by spatial variation, with one corner of the farm showing a lower yield (Figure 7A). In general, the proportion of spatial variance was larger at Lerøy (Figure 7B) than at Austevoll (Figure 7C).

**Figure 7.**
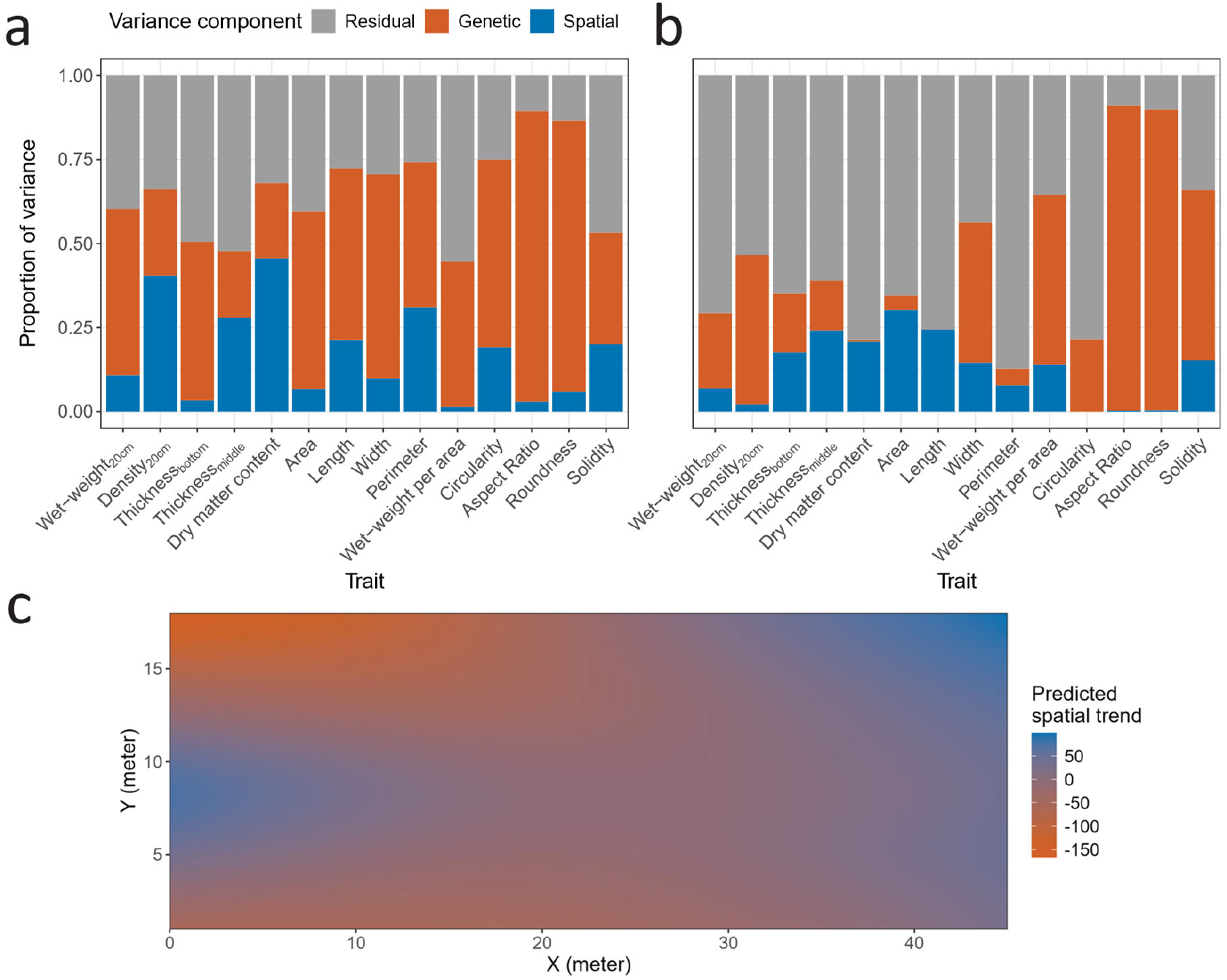
(a) Variance partitioning for Lerøy. (b) Variance partitioning for Austevoll. (c) Spatial trend for wet-weight20cm for Lerøy.

Furthermore, the proportion of genetic variance also differed between the traits and farms, with several traits, such as dry matter content, area, length, and perimeter, showing low genetic variance proportion at Austevoll. These traits should therefore be interpreted with more caution due to unknown environmental or measurement-related factors causing variation. Nevertheless, most traits exhibited substantial genotypic variance in both farms, indicating a genetic basis for trait variation. Additionally, the ability of the model to capture spatial trends for most traits reduced residual variation and therefore improved estimation of the genetic effect.

## 4. Discussion

Breeding for higher yield in *S. latissima* relies on phenotypic variation among genotypes and on understanding how yield is affected by environmental factors. In this study, we captured environmental variation both within farms (spatial trend) and between farms (environment effect and GxE) to improve the estimation of genotype performance. For most traits, a large portion of phenotypic variation was explained by genetic differences. This facilitates breeding as response to selection is proportional to the heritability of traits. The three-fold difference among genotypes for wet-weight_20cm_, with the highest performing genotype producing around 5kg m^-1^, highlights the diversity among the genotype panel. The high generalised heritability of 0.9 for wet-weight_20cm_ further supports the large genetic determination and therefore potential for improvement through breeding. One limitation of this study is the relatively small number of genotypes used to estimate genetic parameters, which should be further explored in larger populations. Screening a larger and more diverse population will improve the estimation of heritability of yield and yield-related traits. In addition, we characterized trait relationships to inform genotype selection. Understanding these relationships is essential for designing effective multi-trait selection strategies in breeding programs.

While yield and density exhibited high heritability values, several morphological traits had low heritability at Austevoll (Table S2). We hypothesize that the higher replication in Lerøy (six versus three in Austevoll) allowed for more precise estimation of heritability of morphological traits. This lower replication in Austevoll likely contributed to the smaller heritability values observed, emphasizing the importance of higher replication in seaweed breeding trials. Similar genotype effects for yield, density and morphological traits have been presented in previous research (Bråtelund et al., 2026; Cohen et al., 2025; Gerard et al., 1987; Gonzalez et al., 2025; Huang et al., 2023). Moreover, we extended our research by including the blade shape traits circularity, aspect ratio, roundness, and solidity. We demonstrated that these traits are strongly genotype dependent, which can be used for selection in early developmental stages and for further examining trait relationships. Together, the substantial phenotypic variation and high heritability for wet-weigth_20cm_, density_20cm_ and morphological traits indicate the potential for trait improvement by breeding.

Almost all traits were not only influenced by the genotype but also by the environment. The traits that differed the most between farms were density_20cm_, thickness_middle_, wet-weight per area and the blade area related traits (area, length, width and perimeter). This suggests that even though the density and blade morphology were affected by the environment, it had a minor effect on yield. Our results are consistent with previous research which documented that *S. latissima* exhibits high phenotypic plasticity in morphological traits (Bruhn et al., 2016; Buck & Buchholz, 2005; Chopin et al., 2004; Diehl & Bischof, 2021; Gerard, 1987; Peteiro & Freire, 2013). Although specific on-farm environmental data was unavailable, several observations suggest that environmental conditions differ between the farms. One difference between farms is the presence of fish aquaculture nearby Lerøy but not Austevoll, and potentially lower currents at Austevoll because of its location in a bay. Both factors may have resulted in higher nutrient fluxes at Lerøy, potentially explaining the higher yield and longer blades with higher wet-weight per area. Interestingly, a higher current at Lerøy would also be expected to increase aspect ratio (length/width), which we did not observe. This may instead be due to the higher density_20cm_ at Austevoll, suggesting that density has a stronger influence on blade shape than current. Overall, our results emphasise the plasticity of *S. latissima* resulting in different morphology and yield among farms.

Genotype by environment interaction (GxE) was significant for almost all traits but contributed relatively little to phenotypic variation. Gonzalez et al. (2025) also reported a significant GxE for yield and density traits, but not for morphology. Based on their reported sums of squares, genotype effects were larger than GxE effects, except for the dry yield. Residual variance was not reported in their study, so these values do not reflect the total variation. The magnitude of GxE effects largely depends on the choice of genotypes and environments. In seaweed farms, some environmental factors, such as temperature, vary over longer distances between farms, while others, such as nutrient concentration and current, can vary strongly even between nearby farms. Gonzalez et al. (2025) observed a larger difference in yield between farms than our study, suggesting that yield may be more sensitive to large environmental differences and that optimal site selection is important to boost yield (Kerrison et al., 2015). Overall, our study showed that GxE has a limited effect, which is promising from a breeding perspective, as genotype performance can be reliably estimated across locations. Nevertheless, additional studies across different environments and genotypes are needed to better understand the role of GxE in *S. latissima* breeding trials.

Environmental differences do not only occur between farms, but also within farms. We demonstrated that accounting for this spatial variation on the farm improved genotype performance estimation. Spatial variation was present on both farms, although the magnitude differed among farms and traits. The lower spatial variation observed at Austevoll may be due to farm-specific characteristics, such as lower environmental heterogeneity due to the location within a bay or an orientation less aligned with environmental gradients. Additionally, at Lerøy we had higher statistical power for morphological traits because of six replicates compared to the standard three replicates at Austevoll. Regardless of the differences among the farms, the consistent presence of spatial variation on both farms stresses the importance of correcting for plot location on the farm for *S. latissima* breeding trials. Only few studies on kelp have corrected for spatial variation. Bråtelund et al. (2026) corrected for spatial variation and suggested that rope number might have influenced the yield of their trial but lacked replication to test this. This underscores the importance of adequate replication in breeding trials to allow for spatial correction.

Investigation of trait relationships can facilitate breeding by determining traits that are easy and rapid to assess. Given that multivariate trait relationships were consistent among genotypes and farms, the associations are likely to be broadly generalisable. Our yield trait, wet-weight_20cm_, was strongly and positively correlated with thickness measurements and the blade size-related traits area, length, and perimeter. Thickness measurements and visual scoring of blade size can therefore serve as a proxy for yield and accelerate breeding. As thickness and blade area are positively correlated, selection for one trait would likely increase the other as well as the yield. The positive relationship further suggests that larger blades may result from an increased cell size or additional cell layers. These traits can be non-destructively assessed throughout the growing season and have the potential to be used in early-stage selection if proven to be correlated with final yield. However, because wet-weight_20cm_ was negatively correlated to dry matter content of individual blades, the relationship between whole plot dry weight, thickness and blade area-related traits should be further investigated.

To our knowledge, this is the first attempt to investigate blade shape traits in relation to yield and biomass allocation over the blade. We did not detect a correlation between yield and the blade shape traits circularity, aspect ratio, roundness, and solidity, suggesting that blade shape has limited value for predicting genotype performance. However, density_20cm_ was correlated with both perimeter and aspect ratio, indicating that higher density results in longer blades with smoother blade edges. Even though the thickness of a single blade in a rope segment was positively correlated with yield, wet-weight per area produced only a weak and non-significant positive correlation with blade thickness. This indicates that these local measurements cannot completely capture variation in thickness across the sporophyte. Thickness_middle_ correlated slightly more strongly with wet-weight per area than thickness_bottom,_ and is thus a better prediction for overall blade thickness. Together, our findings suggest that blade shape traits are not accurate predictors of yield, whereas thickness_middle_ seems informative of biomass allocation over the blade.

Trait correlations observed in our study are largely consistent with previously reported correlations (Bråtelund et al., 2026; Cohen et al., 2025; Gonzalez et al., 2025; Huang et al., 2023). Cohen et al. (2025) observed even higher correlations for some traits, likely due to their more controlled tank environment. Their blades were wider at higher densities, which is in contrast with previous studies. Surprisingly, Huang et al. (2023) found no correlation between blade width and both wet weight and thickness, whereas we detected a positive correlation. Overall, our correlations align with previously reported trait relationships.

Previous farm studies have been conducted with different genetic backgrounds in different locations, making it difficult to compare the studies. Seaweed breeding could therefore benefit from systematic investigations of trait relationships and predictors of yield across developmental stages and environments. Growing the same genotypes in land-based systems and on farms would enable disentangling environmental effects on traits. At the same time, this would allow traits to be measured at earlier developmental stages, enabling early selection and extrapolation of genotype performance from tanks to farm. Genotype selection on land could substantially reduce breeding costs, as farm access is expensive and often limited by harsh environmental conditions. This would allow a drastic increase in the number of genotypes to screen. In addition, phenotyping accuracy may improve because transport of seaweed to land for assessment can affect sporophyte quality due to rapid deterioration after harvest. Farm trials are currently limited in the extend of genetic variation that can be assessed, as the risks of introducing non-local genetic variation into wild populations remain poorly understood. However, screening large allelic variation is essential for breeding programs. Land- and lab-based systems would allow for screening this larger diversity and development of molecular markers by genome wide association studies (GWAS) or quantitative trait loci (QTL) mapping. Moreover, if early growth of sporophytes or even gametophytes correlate with final on-farm yield, lab-based systems could enable efficient screening of large genetic diversity with limited resources.

In this study, we demonstrated that local *S. latissima* genotypes harbour substantial phenotypic variation that can be exploited for breeding for increased yield. At present, breeding programs still rely on local genetic variation to minimise the risk of introducing non-local genetic material into wild populations (Campbell et al., 2019; Goecke et al., 2020). Although the North-West European seaweed farming sector is still considered small, upscaling could increase the ecological risks associated with large-scale cultivation. Strategies such as the production of sporeless sporophytes or gaining a better understanding of population dynamics may enable safe introduction of non-local genetic material in the future (Goecke et al., 2020; Vissers et al., 2024). Nevertheless, our results indicated that considerable variation already exists within local populations, and that this local variation could be used for targeted improvement of yield and its underlying morphological traits.

## Acknowledgements

We would like to thank the companies and owners of the farms where the farm trials were conducted, Lerøy Ocean Harvest and Austevoll Sea Organic AS. We thank Imke Meyer and Joey Visser for their help with phenotyping. We would like to thank Maria João Caldas Paulo for input on experimental design, Martin Boer for input on the spatial analysis and Antonio Lippolis for feedback on the experimental design and manuscript. We thank the SEASEEDS consortium for feedback during the planning and analysis of the experiment.

## Data availability

The data underlying this article will be shared on reasonable request to the corresponding author.

## Declaration of generative AI and AI-assisted technologies in the manuscript preparation process

During the preparation of this work the author(s) used ChatGPT in order to improve writing and R code debugging. After using this tool, the authors reviewed and edited the content as needed and take full responsibility for the content of the published article.

## Data availability

Data will be made available on request.

## CRediT authorship contribution statement

Sanne Put: Writing – original draft, Methodology, Investigation, Formal analysis, Conceptualization, Data curation, Resources, Visualization. Andries Temme: Methodology, Supervision, Writing – review & editing. Jessica Schiller: Investigation, Methodology, Resources. Brigit Reus: Investigation, Methodology, Resources. Gabriel Montecinos Arismendi: Investigation, Methodology, Resources. Tijs Ketelaar: Funding acquisition, Supervision, Writing – review & editing. Luisa M. Trindade: Funding acquisition, Supervision, Writing – review & editing, Conceptualization

## References

Araus, J. L., & Cairns, J. E. (2014). Field high-throughput phenotyping: The new crop breeding frontier. Trends in Plant Science, 19(1), 52–61. 10.1016/j.tplants.2013.09.008

Ball, S. T., Mulla, D. J., & Konzak, C. F. (1993). Spatial heterogeneity affects variety trial interpretation. Crop Science, 33(5), 931–935. 10.2135/cropsci1993.0011183X003300050011X

Boer, M. P. (2023). Tensor product P-splines using a sparse mixed model formulation. Statistical Modelling, 23(5–6), 465–479. 10.1177/1471082X231178591

Bråtelund, S., Ødegård, J., Klemetsdal, G., Ruttink, T., & Ergon. (2026). A quantitative genetic analysis of size-related traits in cultivated sugar kelp (Saccharina latissima). Aquaculture, 611, 743030. 10.1016/J.AQUACULTURE.2025.743030

Bruhn, A., Tørring, D., Thomsen, M., Canal-Vergés, P., Nielsen, M., Rasmussen, M., Eybye, K., Larsen, M., Balsby, T., & Petersen, J. (2016). Impact of environmental conditions on biomass yield, quality, and bio-mitigation capacity of Saccharina latissima. Aquaculture Environment Interactions, 8, 619–636. 10.3354/aei00200

Buck, B. H., & Buchholz, C. M. (2005). Response of offshore cultivated Laminaria saccharina to hydrodynamic forcing in the North Sea. Aquaculture, 250(3–4), 674–691. 10.1016/j.aquaculture.2005.04.062

Campbell, I., Macleod, A., Sahlmann, C., Neves, L., Funderud, J., Øverland, M., Hughes, A. D., & Stanley, M. (2019). The environmental risks associated with the development of seaweed farming in Europe - prioritizing key knowledge gaps. Frontiers in Marine Science, 6. 10.3389/fmars.2019.00107

Chopin, T., Robinson, S., Sawhney, M., Bastarache, S., Belyea, L., Shea, R., Armstrong, W., Stewart, I., & Fitzgerald, P. (2004). The AquaNet integrated multi-trophic aquaculture project: Rationale of the project and development of kelp cultivation as the inorganic extractive component of the system. Bulletin of the Aquaculture Association of Canada, 104(3), 11–18.

Cohen, J., Twijnstra, R., Schiller, J., Montecinos Arismendi, G., Reus, B., Soetaert, K., & Timmermans, K. (2025). Global interfertility and heterosis in sugar kelp populations: a next step in sugar kelp breeding. Journal of Applied Phycology, 37(2), 1213–1226. 10.1007/s10811-025-03447-7

Diehl, N., & Bischof, K. (2021). Coping with a changing Arctic. Source: Marine Ecology Progress Series, 657, 43–57. 10.2307/26988393

Ebbing, A. P. J., Fivash, G. S., Martin, N. B., Pierik, R., Bouma, T. J., Kromkamp, J. C., & Timmermans, K. (2021). In-culture selection and the potential effects of changing sex ratios on the reproductive success of multiannual delayed gametophytes of saccharina latissima and alaria esculenta. Journal of Marine Science and Engineering, 9(11). 10.3390/jmse9111250

EC. (2022). Commission from the Commission: Towards a Strong and Sustainable EU Algae Sector.

Fernand, F., Israel, A., Skjermo, J., Wichard, T., Timmermans, K. R., & Golberg, A. (2017). Offshore macroalgae biomass for bioenergy production: Environmental aspects, technological achievements and challenges. In Renewable and Sustainable Energy Reviews (Vol. 75, pp. 35–45). Elsevier Ltd. 10.1016/j.rser.2016.10.046

Gerard, V. A. (1987). Hydrodynamic streamlining of Laminaria saccharina Lamour. in response to mechanical stress. Journal of Experimental Marine Biology and Ecology, 107(3), 237–244. 10.1016/0022-0981(87)90040-2

Gerard, V. A., Dubois, K., & Greene, R. (1987). Growth responses of two Laminaria saccharina populations to environmental variation. In Hydrobiologia (Vol. 1511152).

Goecke, F., Klemetsdal, G., & Ergon, Å. (2020). Cultivar Development of Kelps for Commercial Cultivation—Past Lessons and Future Prospects. Frontiers in Marine Science, 8. 10.3389/fmars.2020.00110

Gonzalez, S. T., Li, Y., Aydlett, M., Bailey, D., Kerr, H., Doall, M., Gobler, C. J., Chambers, M., Jannink, J. L., Yarish, C., & Lindell, S. (2025). Evaluation of six sugar kelp crosses selected for high yield at three Northeastern US farms. Aquaculture, 600, 742191. 10.1016/J.AQUACULTURE.2025.742191

Hu, Z. M., Shan, T. F., Zhang, J., Zhang, Q. S., Critchley, A. T., Choi, H. G., Yotsukura, N., Liu, F. L., & Duan, D. L. (2021). Kelp aquaculture in China: a retrospective and future prospects. In Reviews in Aquaculture (Vol. 13, Number 3, pp. 1324–1351). John Wiley and Sons Inc. 10.1111/raq.12524

Huang, M., Robbins, K. R., Li, Y., Umanzor, S., Marty-Rivera, M., Bailey, D., Aydlett, M., Schmutz, J., Grimwood, J., Yarish, C., Lindell, S., & Jannink, J. L. (2023). Genomic selection in algae with biphasic lifecycles: A Saccharina latissima (sugar kelp) case study. Frontiers in Marine Science, 10. 10.3389/fmars.2023.1040979

Huang, M., Robbins, K. R., Li, Y., Umanzor, S., Marty-Rivera, M., Bailey, D., Yarish, C., Lindell, S., & Jannink, J. L. (2022). Simulation of sugar kelp (Saccharina latissima) breeding guided by practices to accelerate genetic gains. G3: Genes, Genomes, Genetics, 12(3). 10.1093/g3journal/jkac003

Iñiguez, C., Carmona, R., Lorenzo, M. R., Niell, F. X., Wiencke, C., & Gordillo, F. J. L. (2016). Increased temperature, rather than elevated CO2, modulates the carbon assimilation of the Arctic kelps Saccharina latissima and Laminaria solidungula. Marine Biology, 163(12). 10.1007/s00227-016-3024-6

Kanda, T. (1936). On the Gametophytes of Some Japanese Species of Laminariales. Scientific Papers of the Institute of Algological Research, Faculty of Science, Hokkaido Imperial University, 1(2), 221–260. https://hdl.handle.net/2115/48057

Kerrison, P. D., Stanley, M. S., Edwards, M. D., Black, K. D., & Hughes, A. D. (2015). The cultivation of European kelp for bioenergy: Site and species selection. Biomass and Bioenergy, 80, 229–242. 10.1016/j.biombioe.2015.04.035

Kraan, S. (2013). Mass-cultivation of carbohydrate rich macroalgae, a possible solution for sustainable biofuel production. Mitigation and Adaptation Strategies for Global Change, 18(1), 27–46. 10.1007/s11027-010-9275-5

Li, D., Zhou, Z.-G., Liu, H., & Wu, C. (1999). A new method of Laminaria japonica strain selection and sporeling raising by the use of gametophyte clones. Hydrobiologia, 398, 473–476.

Li, X., Cong, Y., Yang, G., Shi, Y., Qu, S., Li, Z., Wang, G., Zhang, Z., Luo, S., Dai, H., Xie, J., Jiang, G., Liu, J., & Wang, T. (2007). Trait evaluation and trial cultivation of Dongfang No. 2, the hybrid of a male gametophyte clone of Laminaria longissima (Laminariales, Phaeophyta) and a female one of L. japonica. Journal of Applied Phycology, 19(2), 139–151. 10.1007/s10811-006-9120-0

Oakey, H., Verbyla, A., Pitchford, W., Cullis, B., & Kuchel, H. (2006). Joint modeling of additive and non-additive genetic line effects in single field trials. Theoretical and Applied Genetics, 113(5), 809–819. 10.1007/s00122-006-0333-z

Peteiro, C., & Freire, Ó. (2013). Biomass yield and morphological features of the seaweed Saccharina latissima cultivated at two different sites in a coastal bay in the Atlantic coast of Spain. Journal of Applied Phycology, 25(1), 205–213. 10.1007/s10811-012-9854-9

Robinson, N., Winberg, P., & Kirkendale, L. (2013). Genetic improvement of macroalgae: Status to date and needs for the future. Journal of Applied Phycology, 25(3), 703–716. 10.1007/s10811-012-9950-x

Rodríguez-Álvarez, M. X., Boer, M. P., van Eeuwijk, F. A., & Eilers, P. H. C. (2018). Correcting for spatial heterogeneity in plant breeding experiments with P-splines. Spatial Statistics, 23, 52–71. 10.1016/j.spasta.2017.10.003

Sanderson, J. C., Cromey, C. J., Dring, M. J., & Kelly, M. S. (2008). Distribution of nutrients for seaweed cultivation around salmon cages at farm sites in north-west Scotland. Aquaculture, 278(1–4), 60–68. 10.1016/j.aquaculture.2008.03.027

Schindelin, J., Arganda-Carreras, I., Frise, E., Kaynig, V., Longair, M., Pietzsch, T., Preibisch, S., Rueden, C., Saalfeld, S., Schmid, B., Tinevez, J.-Y., White, D. J., Hartenstein, V., Eliceiri, K., Tomancak, P., & Cardona, A. (2012). Fiji: an open-source platform for biological-image analysis. Nature Methods, 9(7), 676–682. 10.1038/nmeth.2019

Šidák, Z. (1967). Rectangular Confidence Regions for the Means of Multivariate Normal Distributions. Source: Journal of the American Statistical Association, 62(318), 626–633.

Simpson, E. H. (1951). The Interpretation of Interaction in Contingency Tables. Source: Journal of the Royal Statistical Society. Series B (Methodological), 13(2), 238–241. https://doi.org/https://www.jstor.org/stable/2984065

Spearman, C. (1904). The proof and measurement of association between two things. The American Journal of Psychology, 15(1), 72–101.

Theodorou, I., Opsahl-Sorteberg, H.-G., & Charrier, B. (2021). Preparation of Zygotes and Embryos of the Kelp Saccharina latissima for Cell Biology Approaches. BIO-PROTOCOL, 11(16). 10.21769/bioprotoc.4132

Tukey, J. W. (1949). Comparing Individual Means in the Analysis of Variance. 5(2), 99–114. https://www.jstor.org/stable/3001913

van den Burg, S. W. K., van Duijn, A. P., Bartelings, H., van Krimpen, M. M., & Poelman, M. (2016). The economic feasibility of seaweed production in the North Sea. Aquaculture Economics and Management, 20(3), 235–252. 10.1080/13657305.2016.1177859

Van Der Molen, J., Ruardij, P., Mooney, K., Kerrison, P., O’Connor, N. E., Gorman, E., Timmermans, K., Wright, S., Kelly, M., Hughes, A. D., & Capuzzo, E. (2018). Modelling potential production of macroalgae farms in UK and Dutch coastal waters. Biogeosciences, 15(4), 1123–1147. 10.5194/bg-15-1123-2018

Vilmin, V., & Van Duren, L. (2021). Modelling seaweed cultivation on the Dutch continental shelf.

Vissers, C., Lindell, S. R., Nuzhdin, S. V., Almada, A. A., & Timmermans, K. (2024). Using sporeless sporophytes as a next step towards upscaling offshore kelp cultivation. Journal of Applied Phycology, 36(1), 313–320. 10.1007/s10811-023-03123-8

Wang, X., Yao, J., Zhang, J., & Duan, D. (2020). Status of genetic studies and breeding of Saccharina japonica in China. Journal of Oceanology and Limnology, 38(4), 1064–1079. 10.1007/s00343-020-0070-1

Warton, D. I., & Hui, F. K. C. (2011). The arcsine is asinine: the analysis of proportions in ecology. Ecology, 92(1), 3–10.

Zhang, Q. S., Tang, X. X., Cong, Y. Z., Qu, S. C., Luo, S. J., & Yang, G. P. (2007). Breeding of an elite Laminaria variety 90-1 through inter-specific gametophyte crossing. Journal of Applied Phycology, 19(4), 303–311. 10.1007/s10811-006-9137-4

