## Supplementary material for "Genotype and farm effects on yield and morphology reveal potential for breeding and site selection for sugar kelp": Figure S1

### Leroy

#### Genotype 1

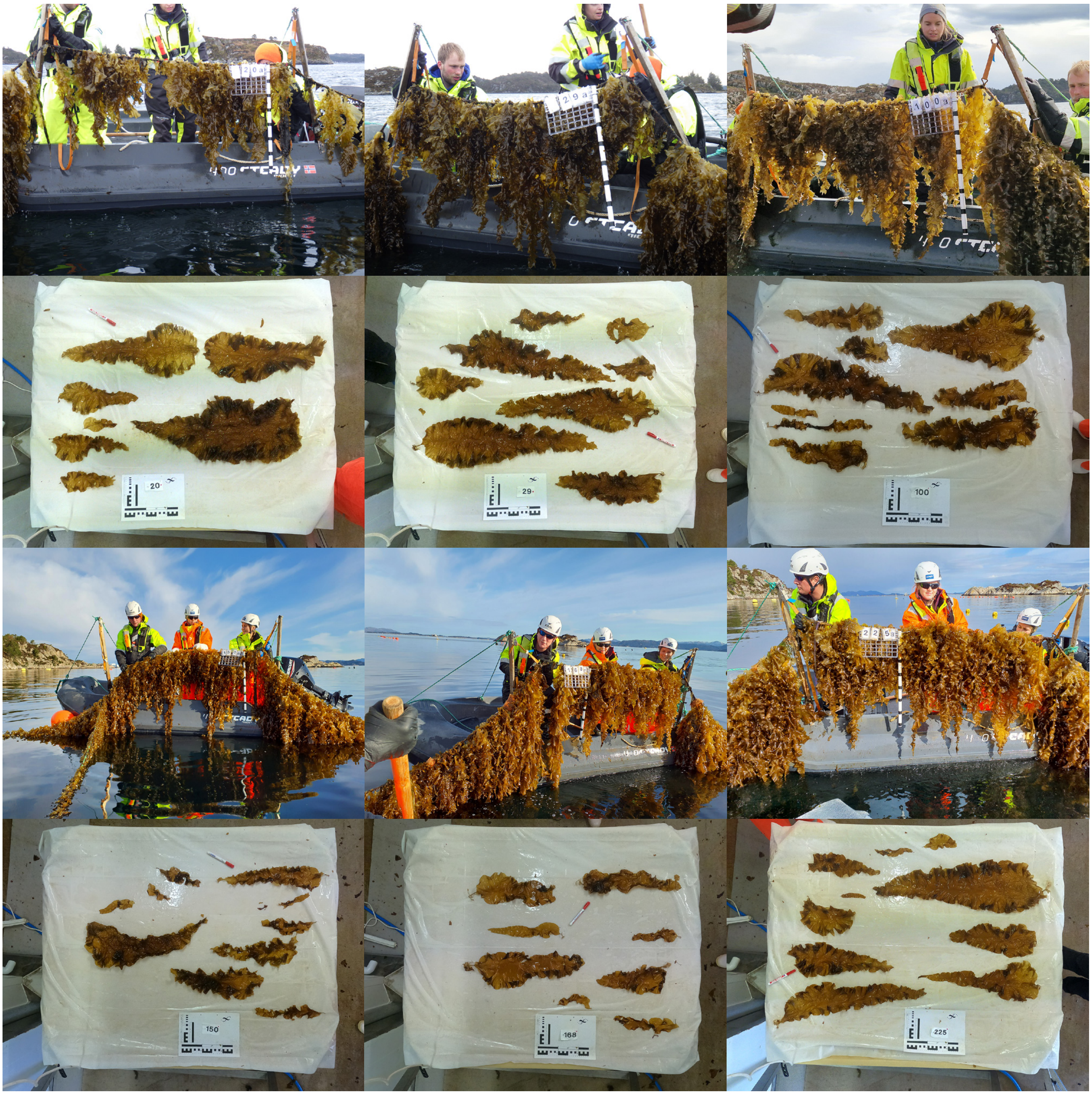


#### Genotype 2

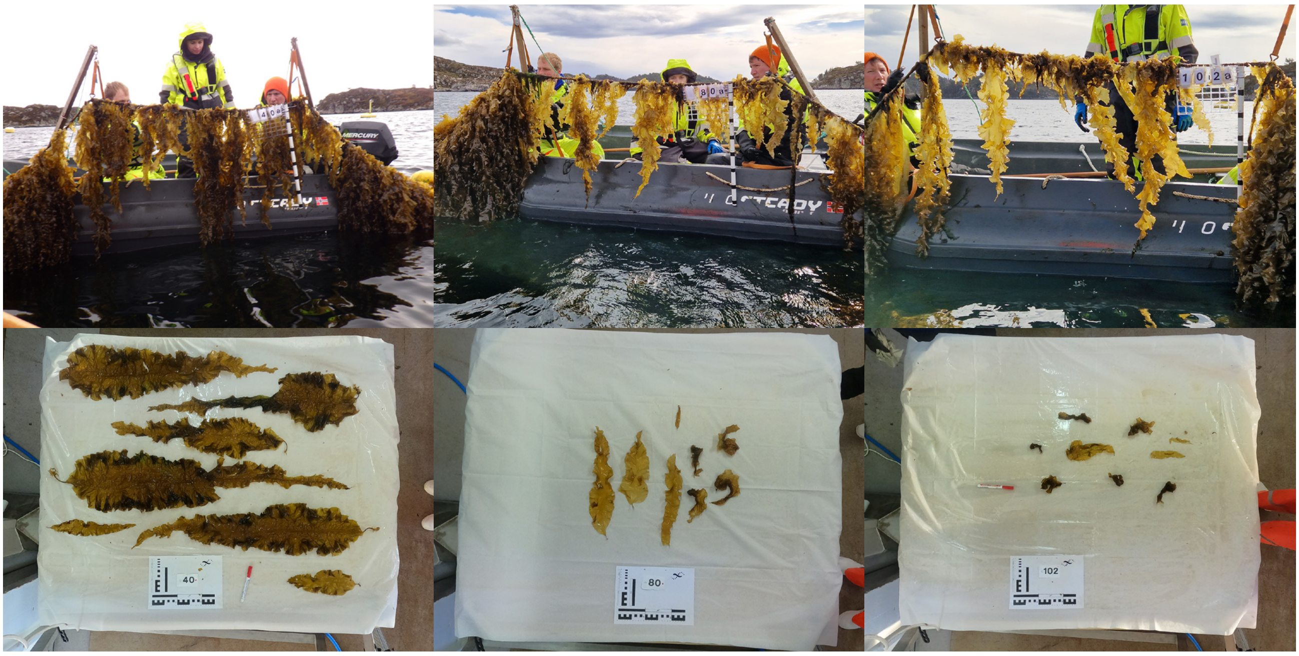

Genotype 2

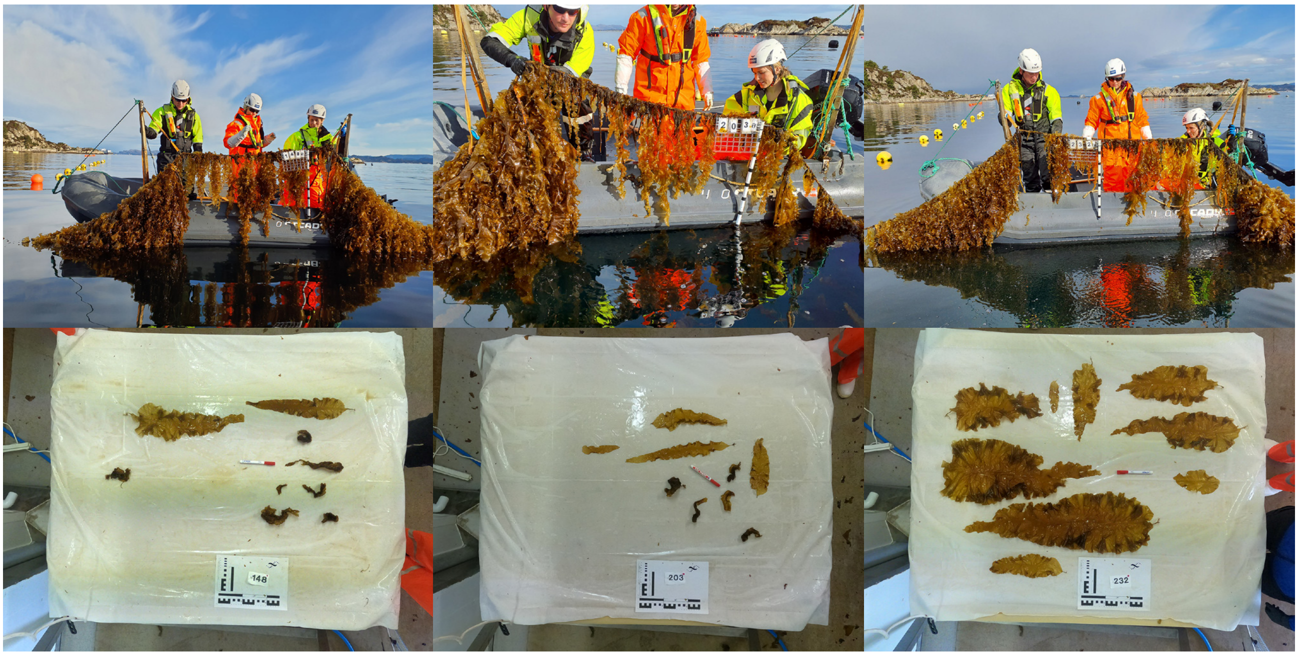


Genotype 3

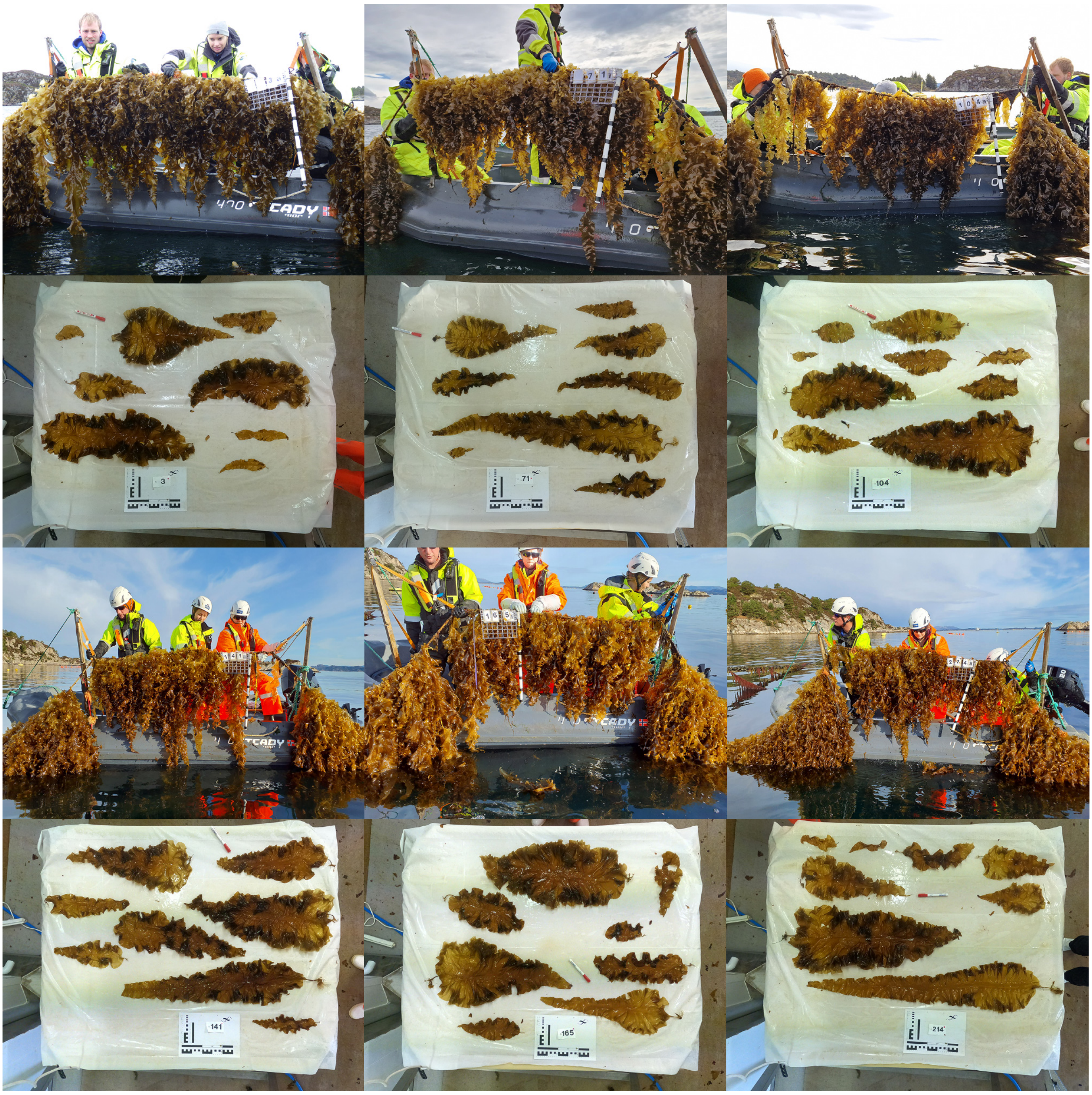

Genotype 4

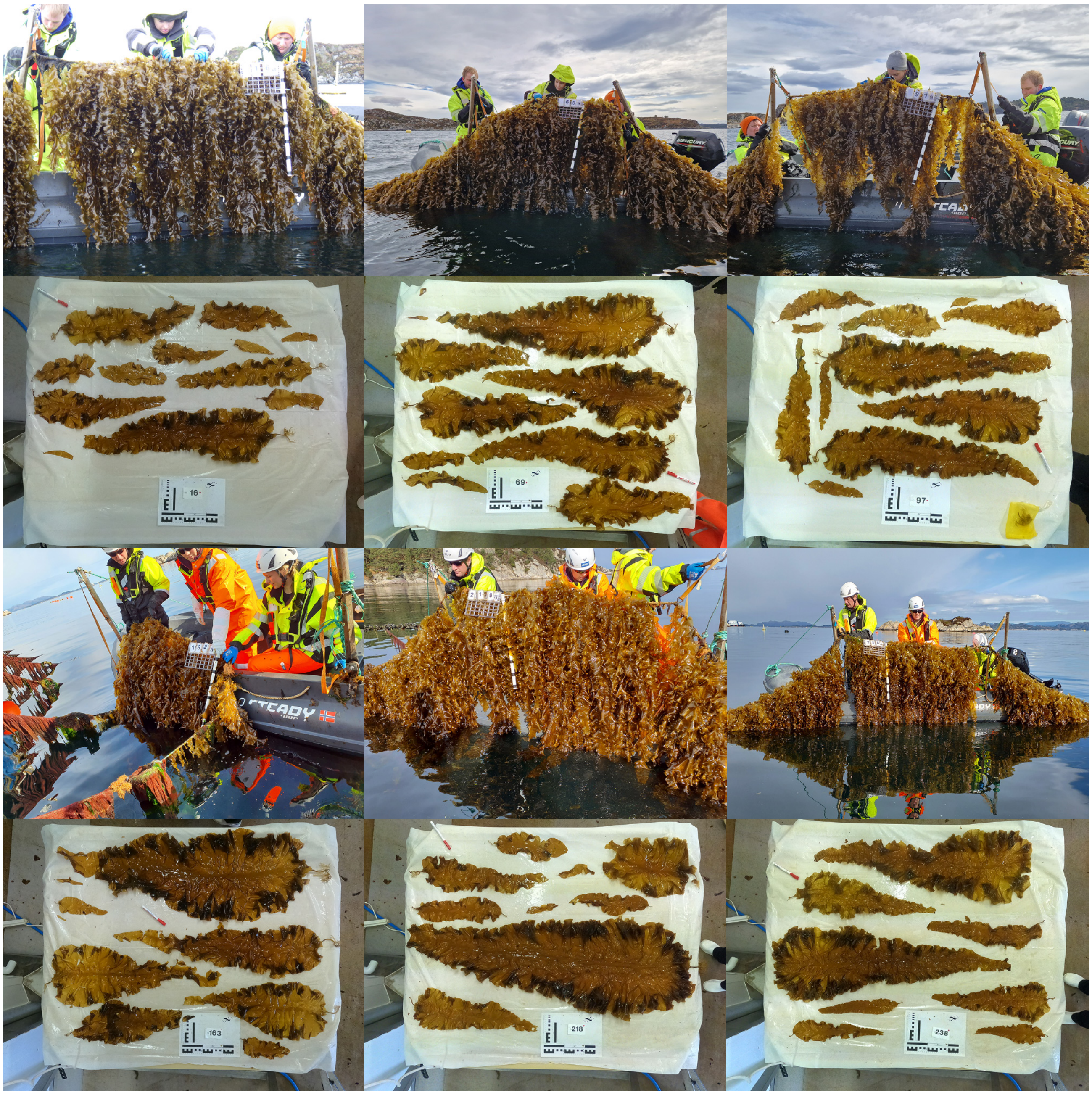

Genotype 5

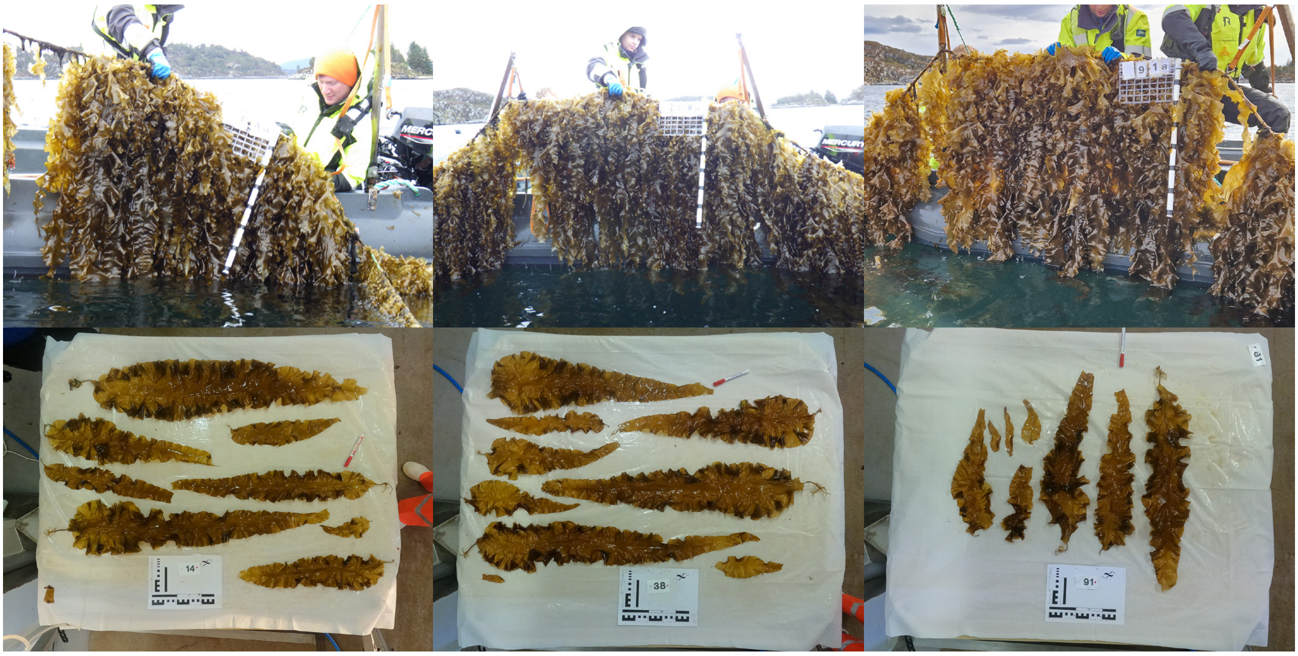

Genotype 5

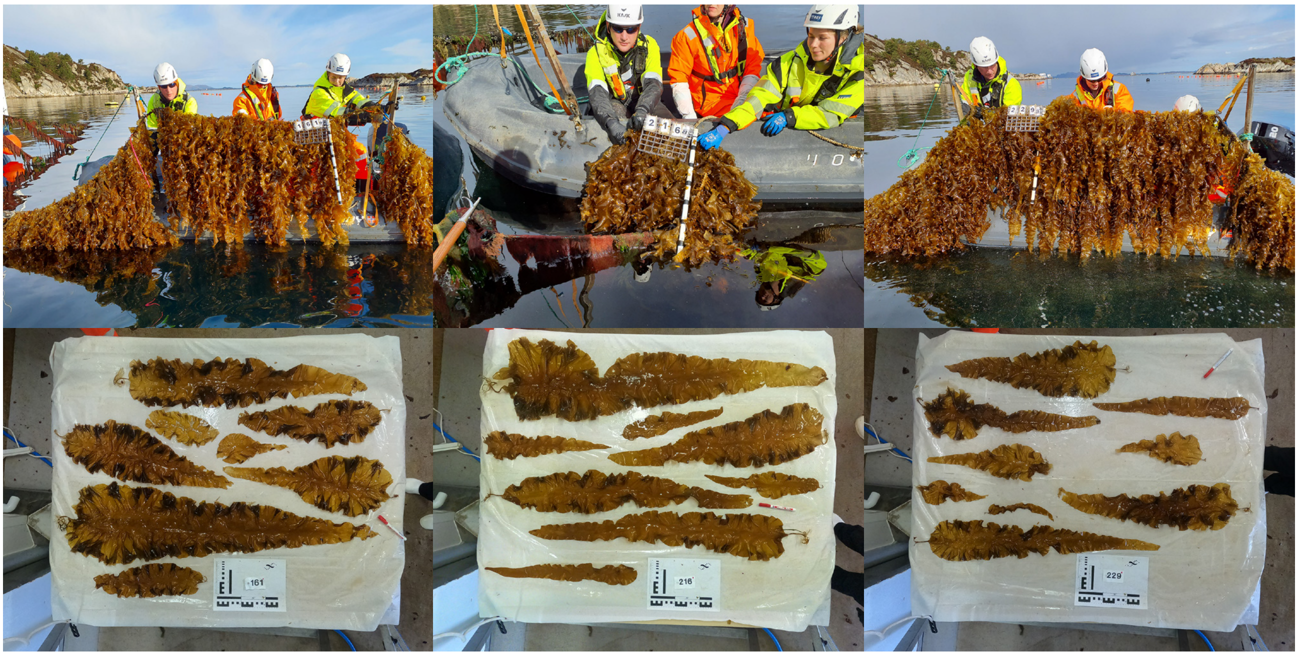


Genotype 6

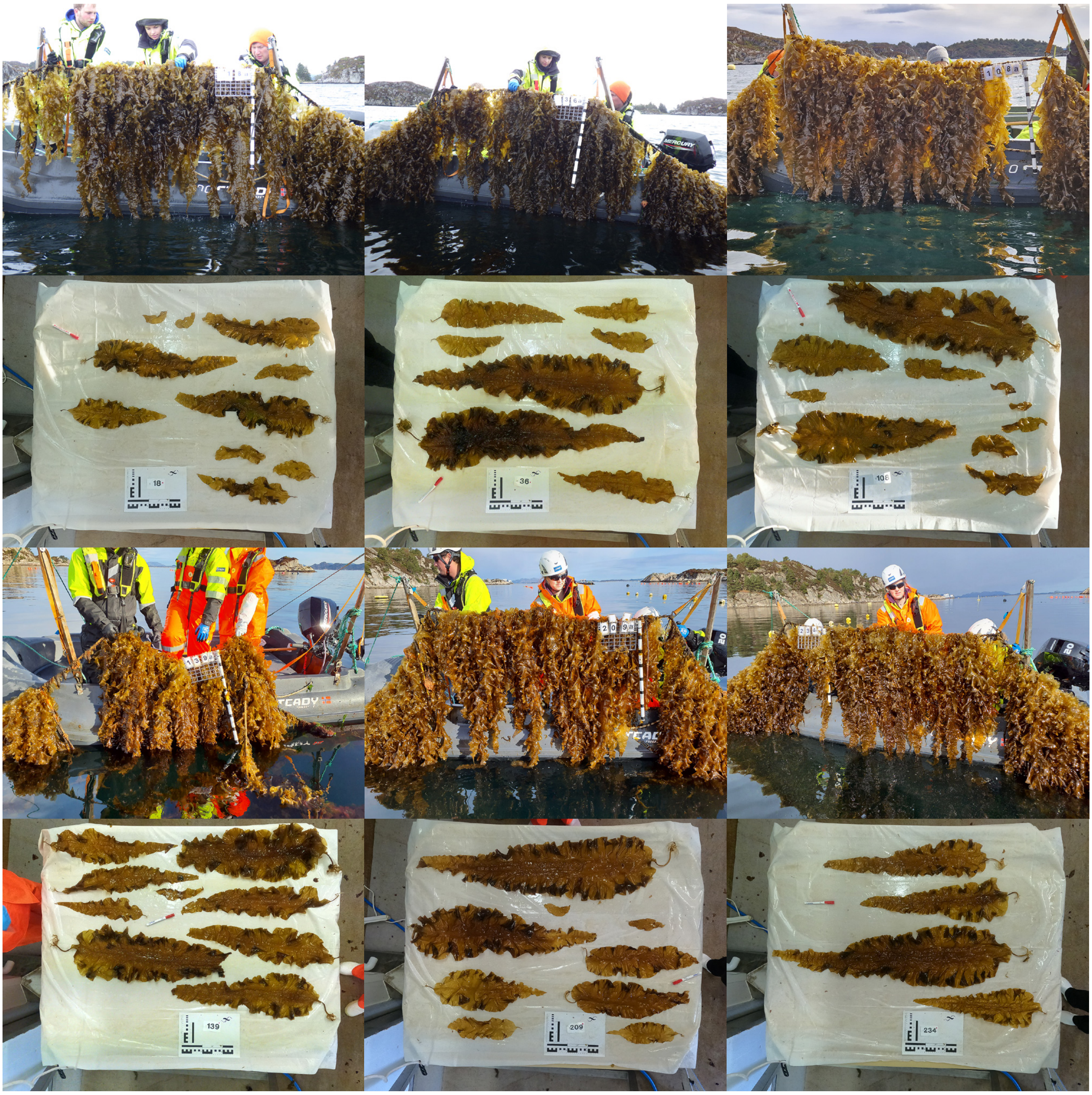

#### Genotype 7

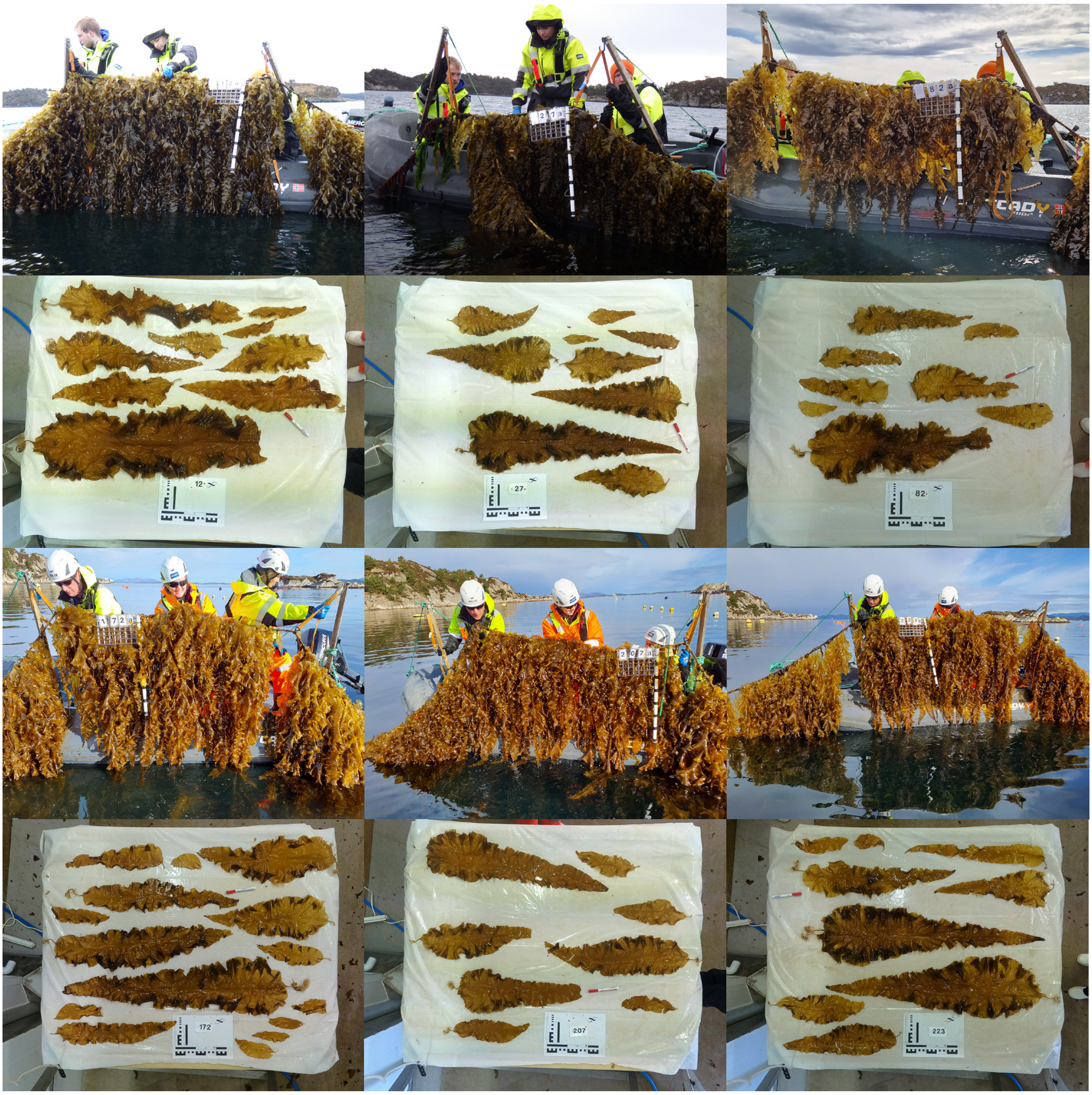


#### Genotype 8

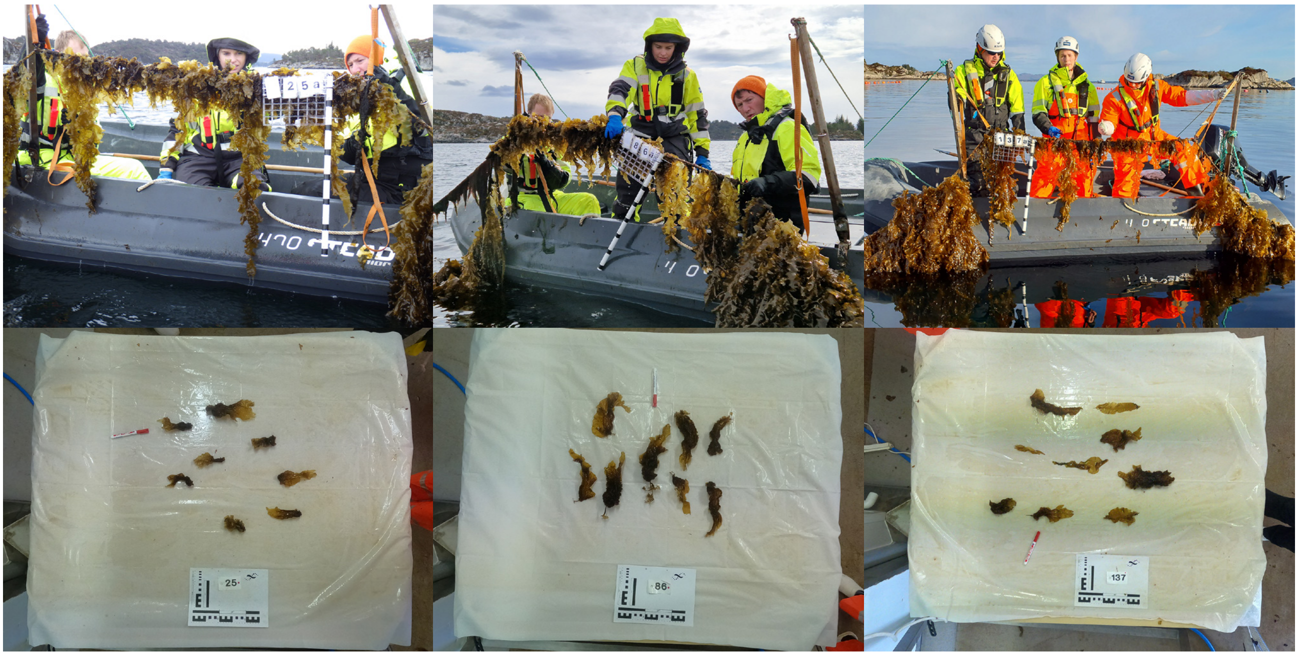

#### Genotype 8

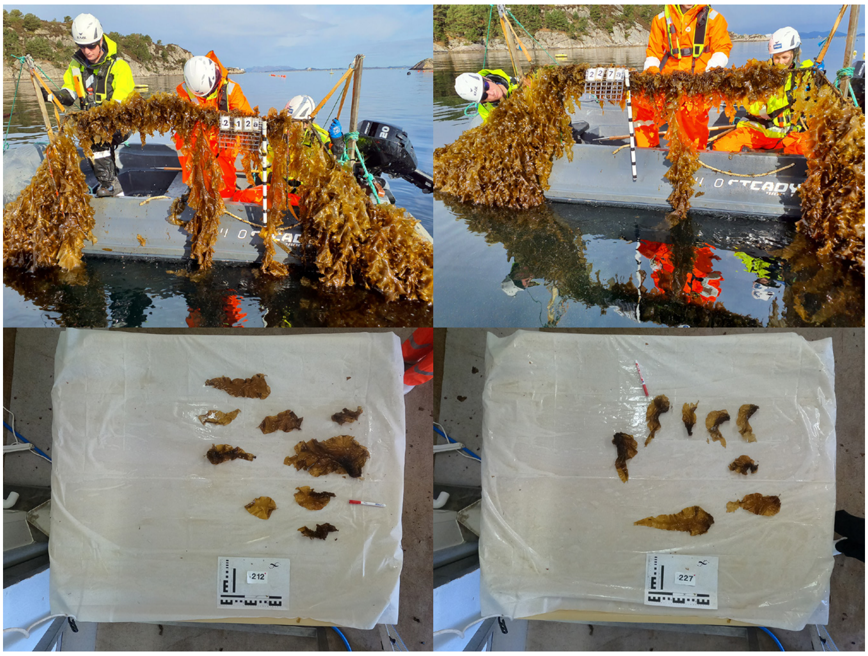


#### Genotype 9

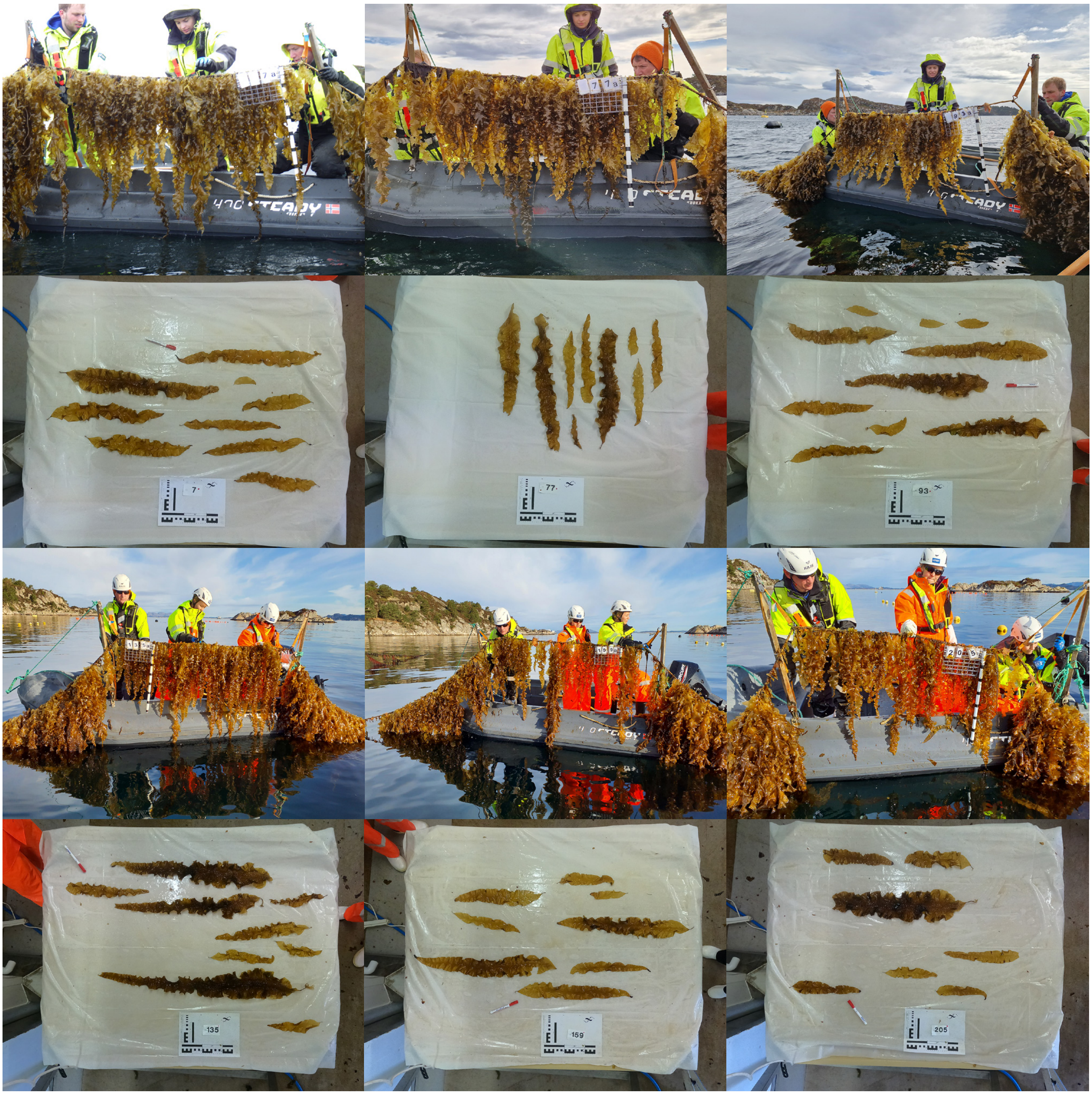

#### Genotype 9

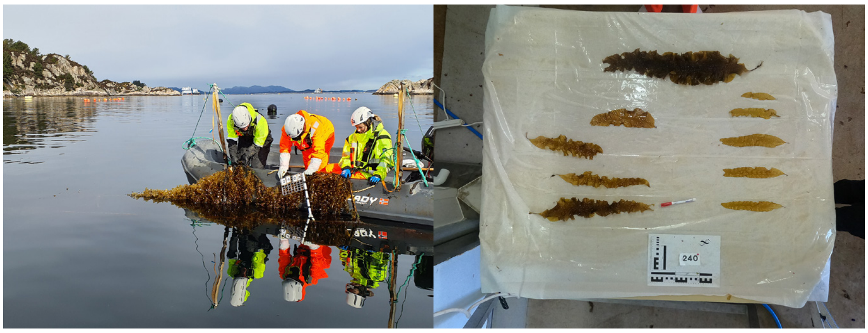


#### Genotype 10

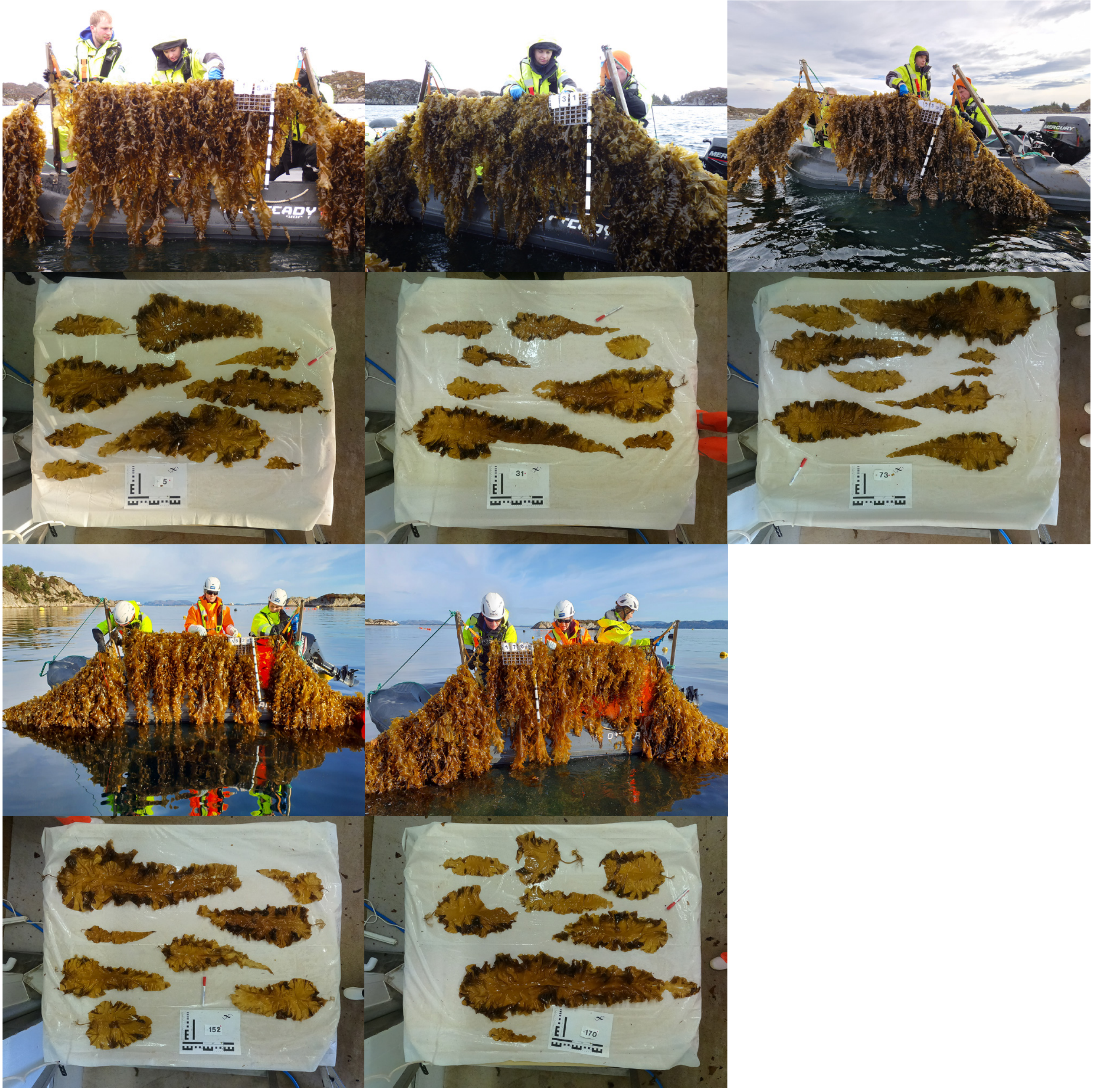

Genotype 11

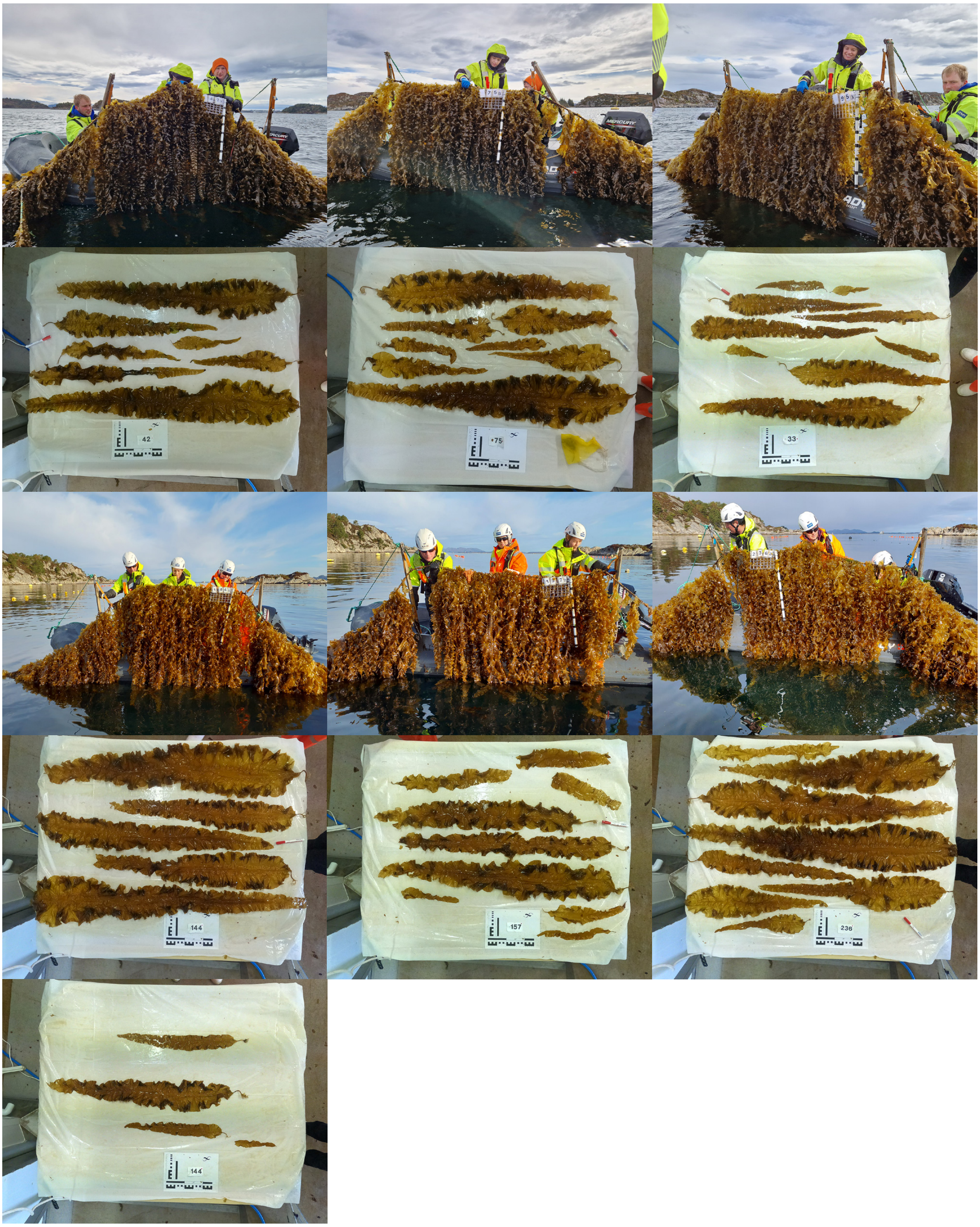

Genotype 12

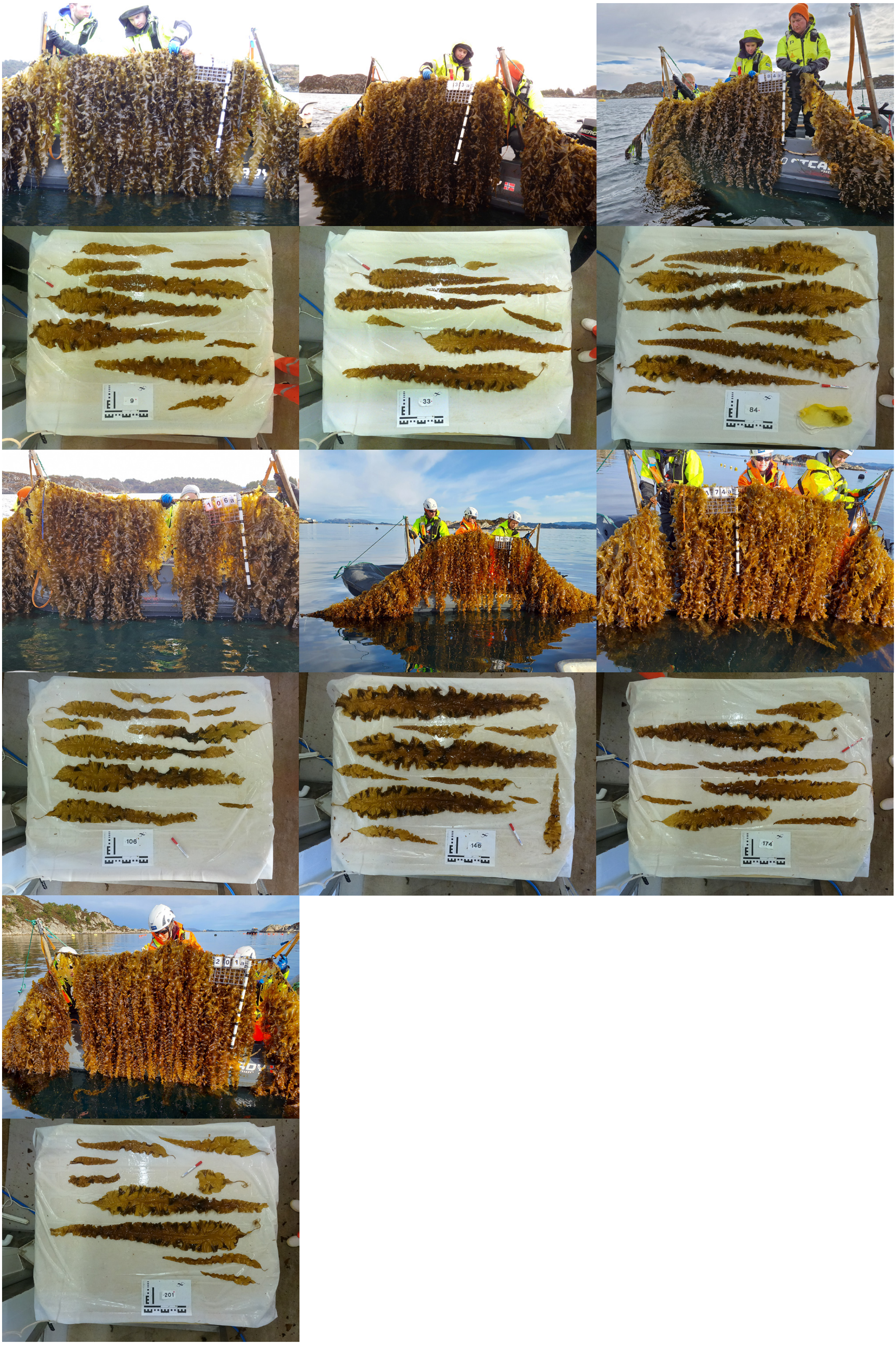

### Austevoll

#### Genotype 1

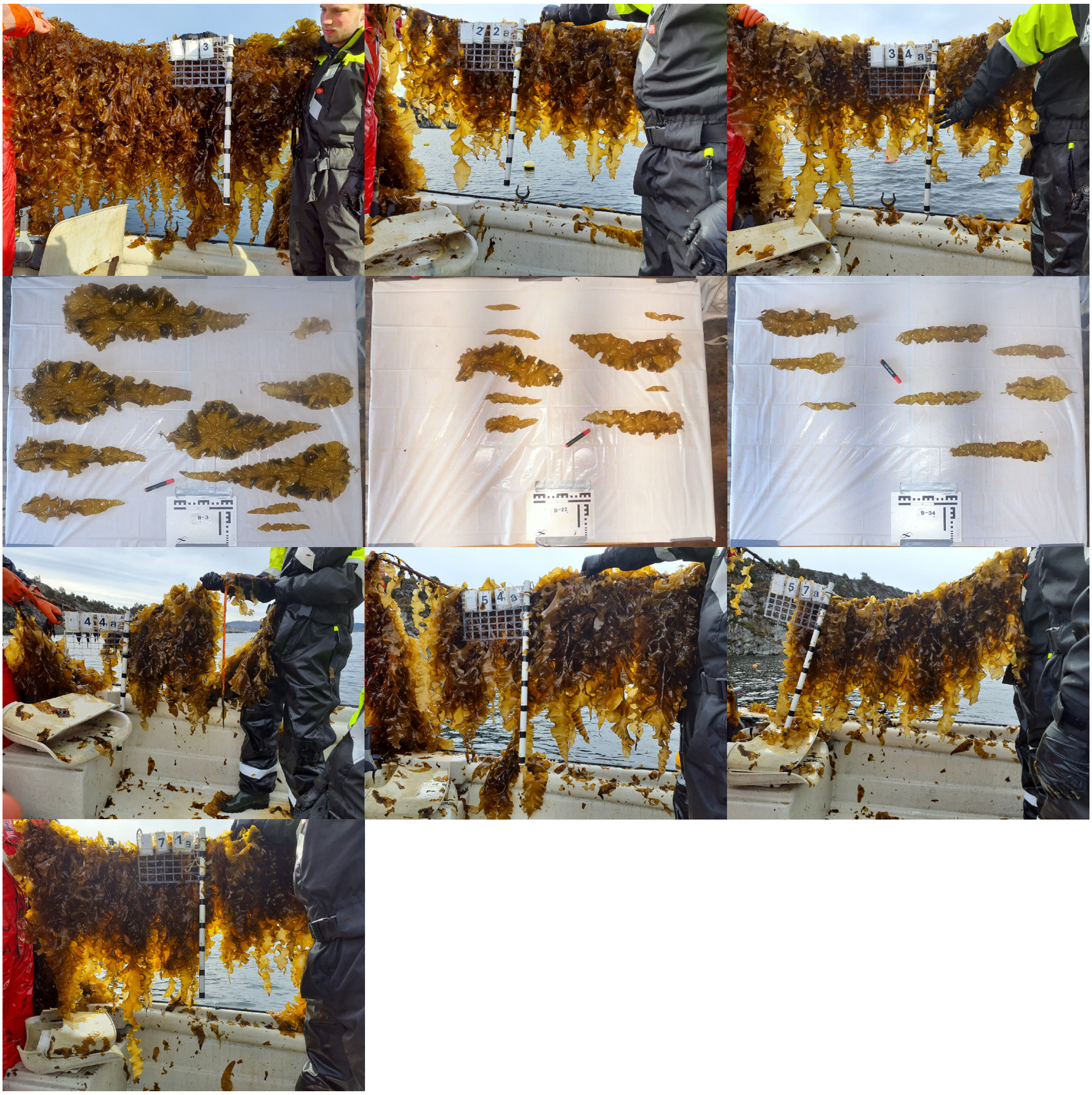

#### Genotype 2

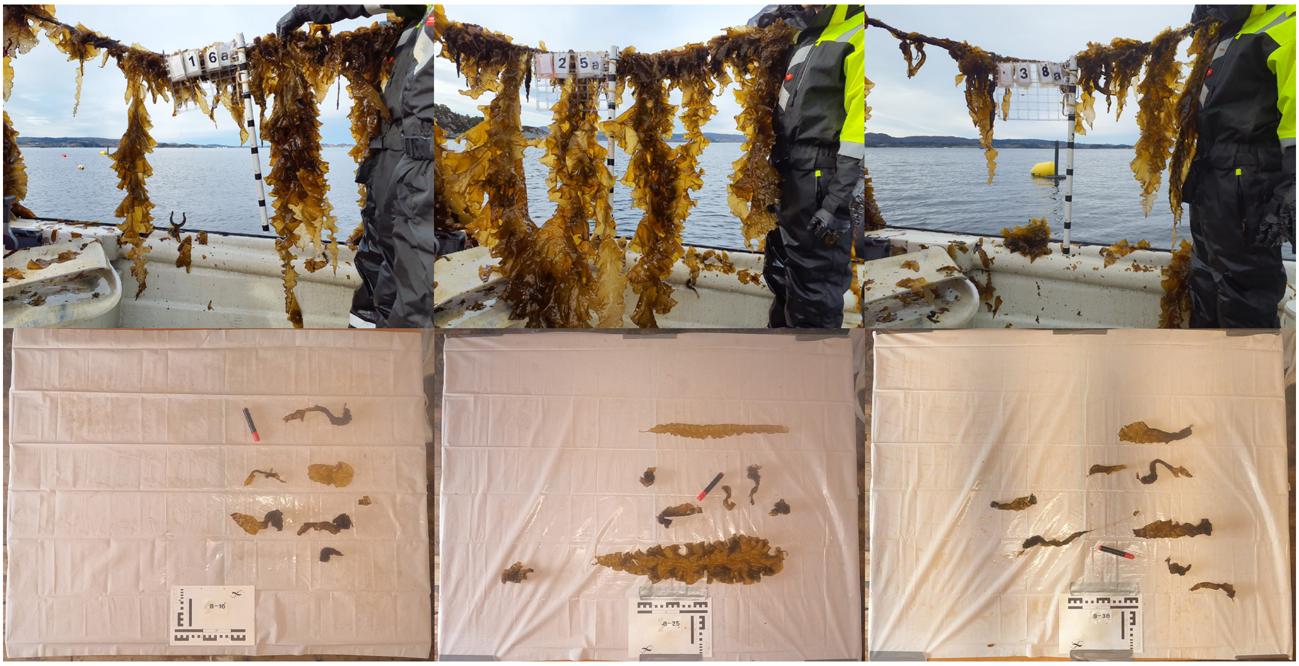

Genotype 2

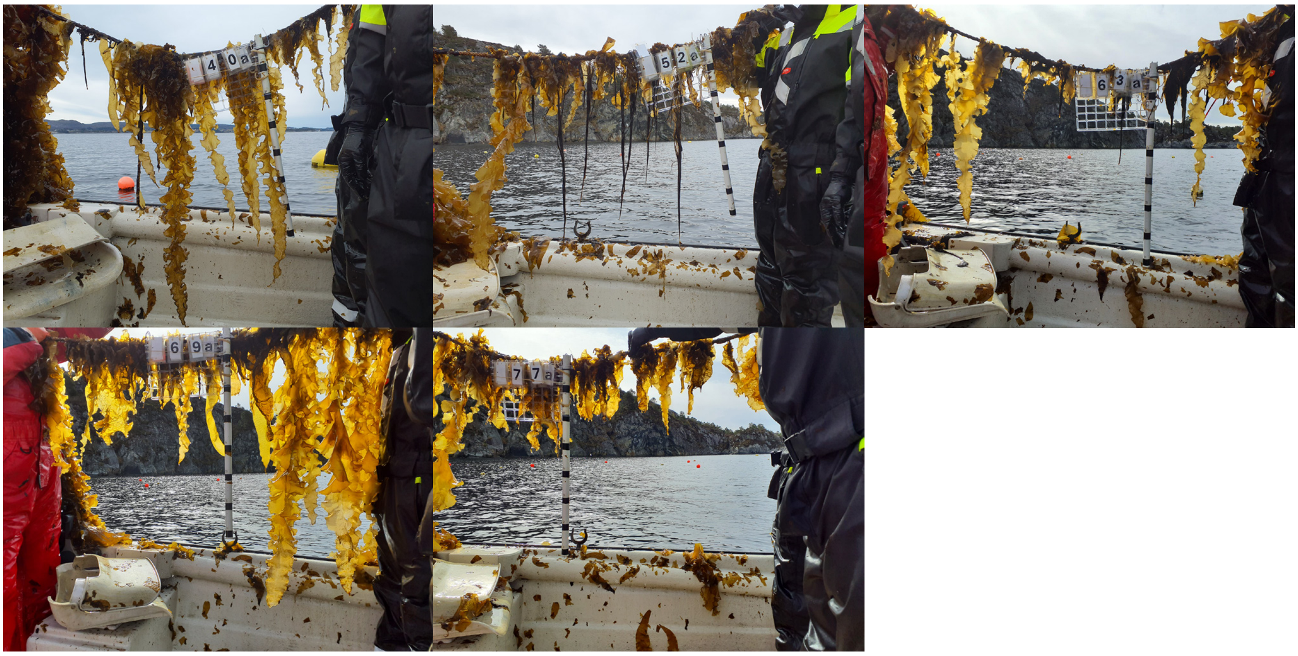


Genotype 3

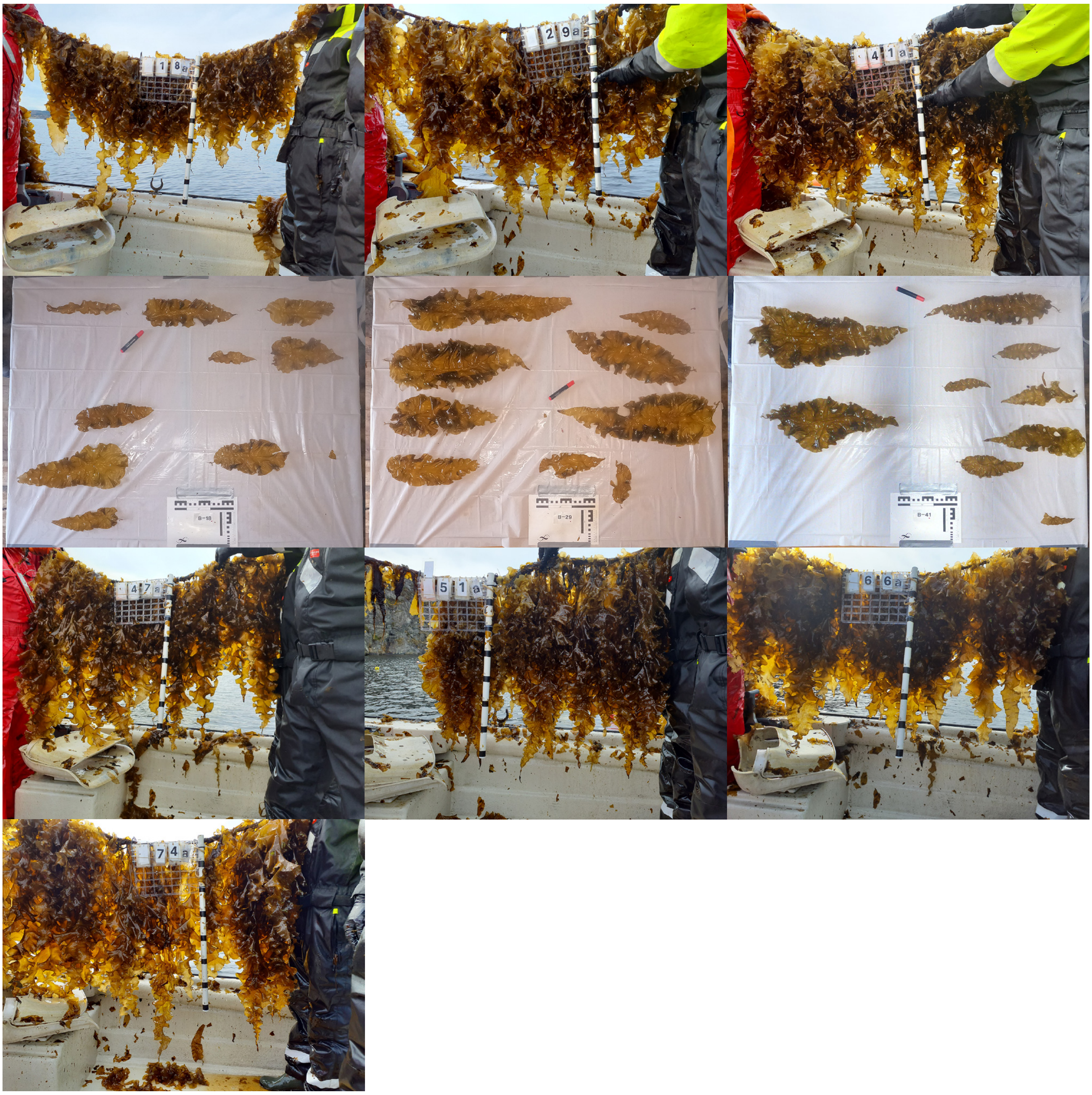

Genotype 4

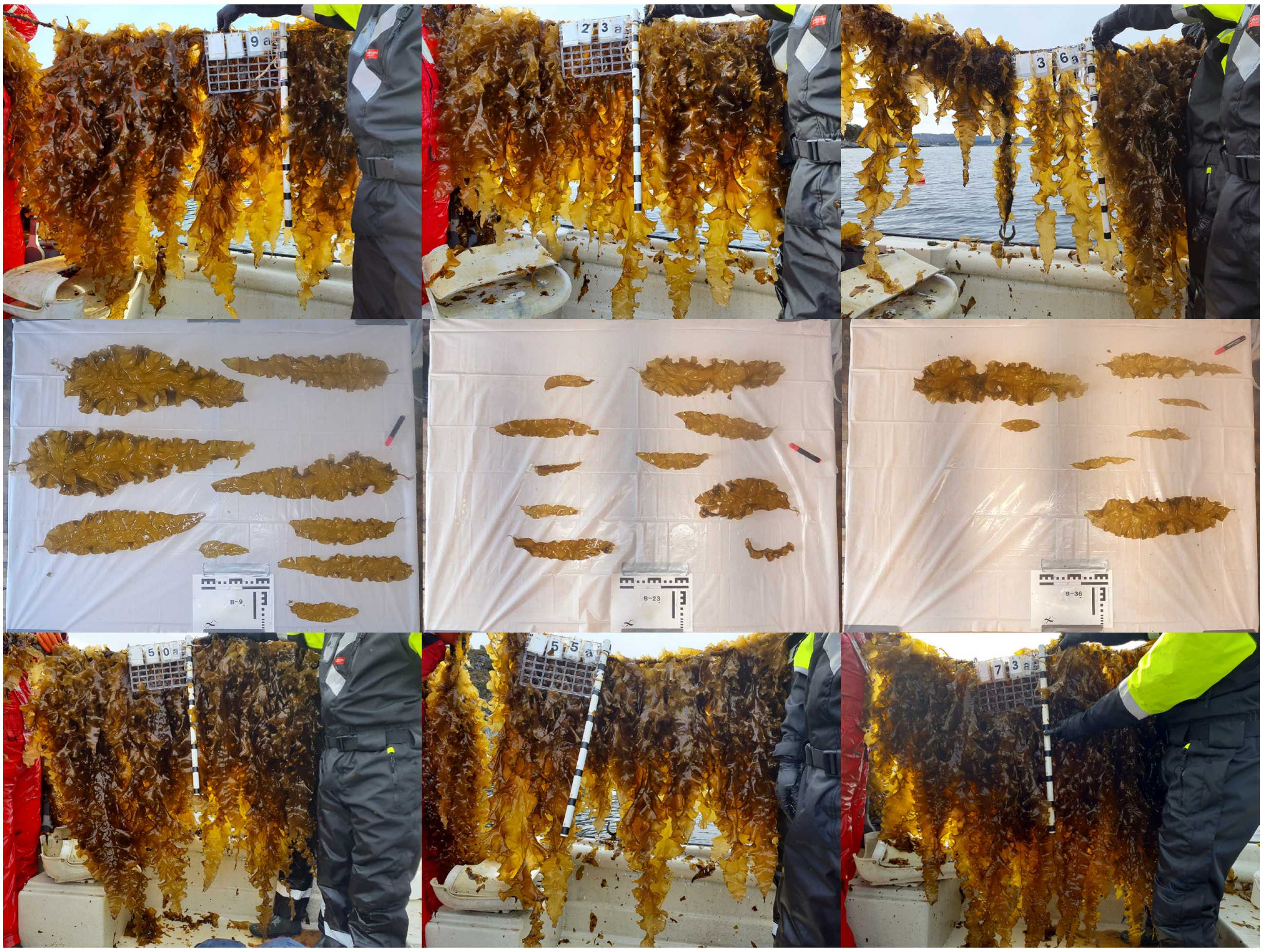

Genotype 5

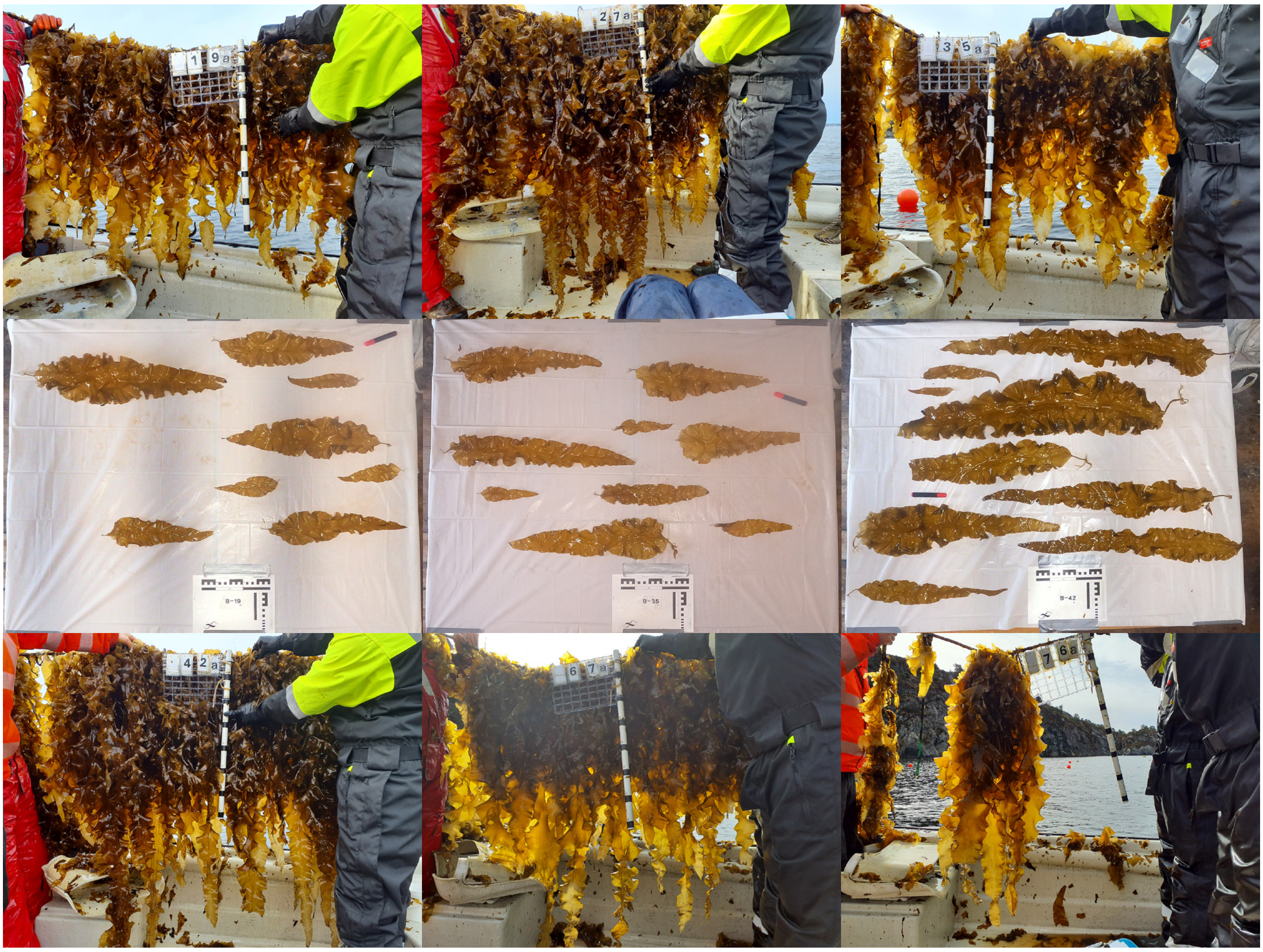

#### Genotype 6

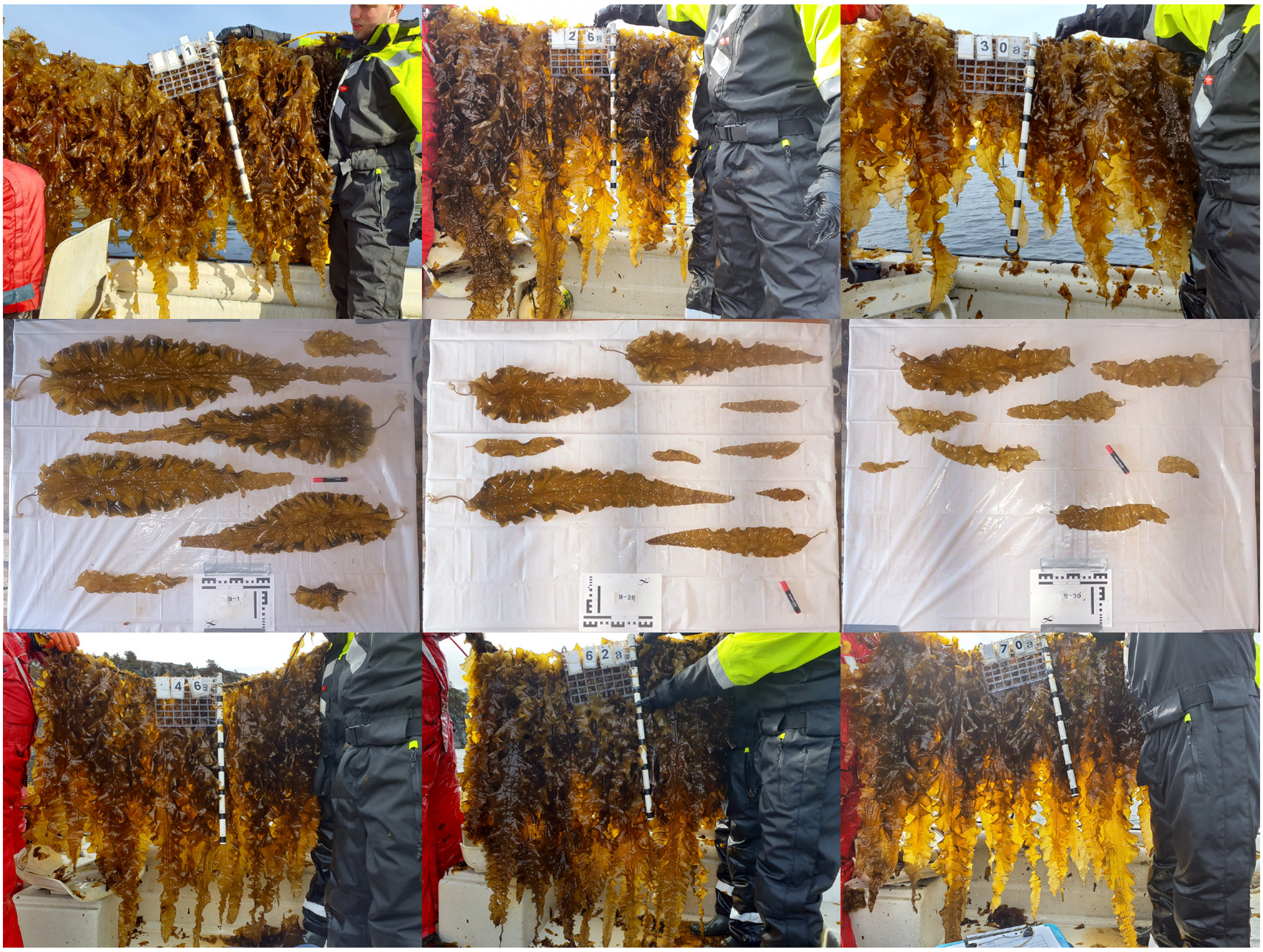

#### Genotype 7

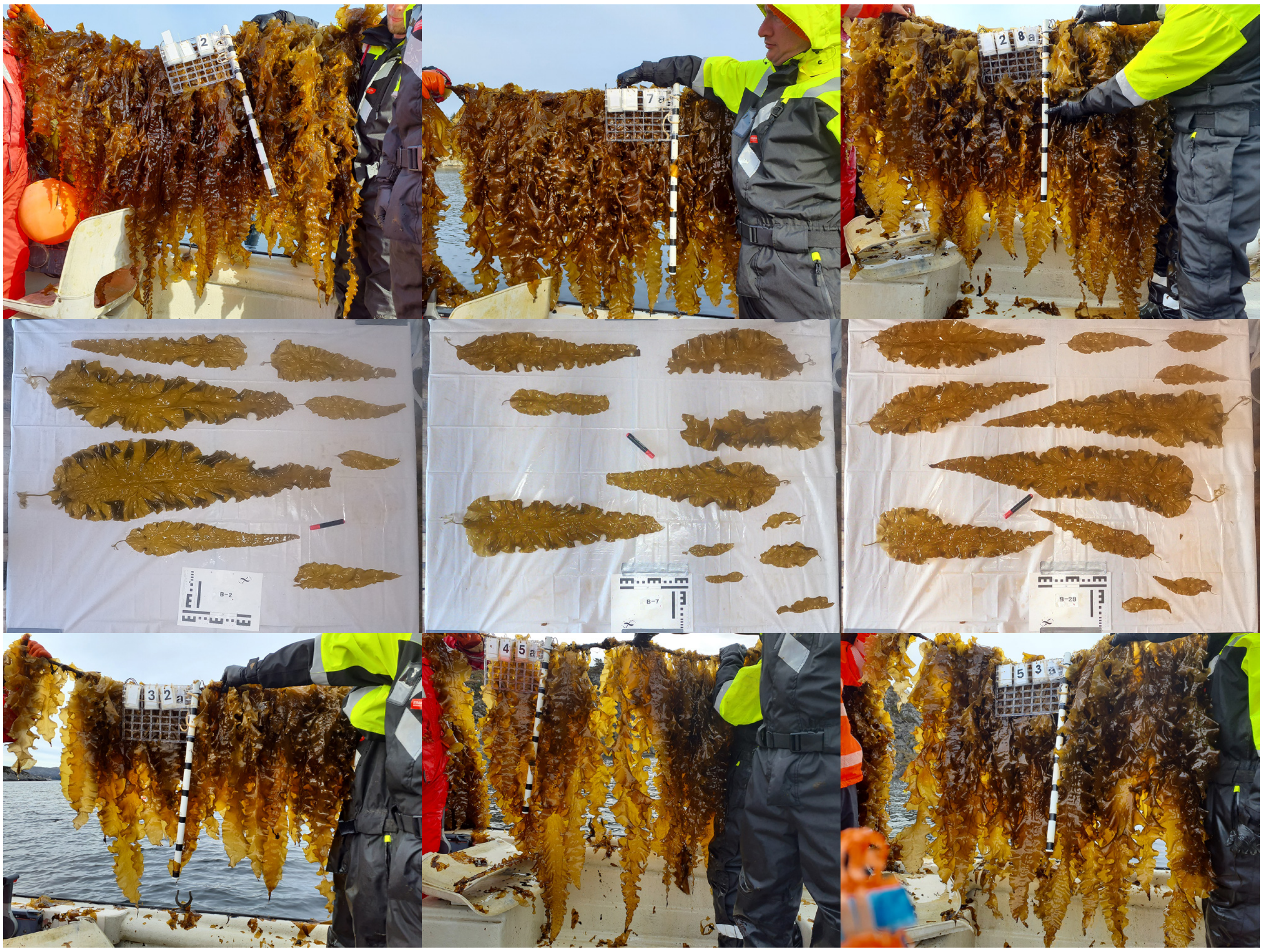

Genotype 7

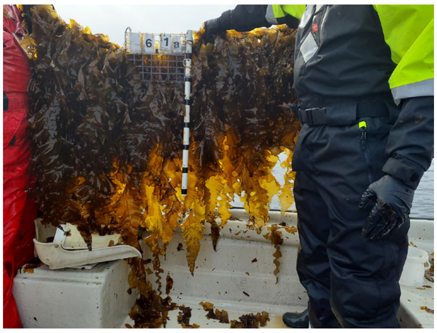


Genotype 8

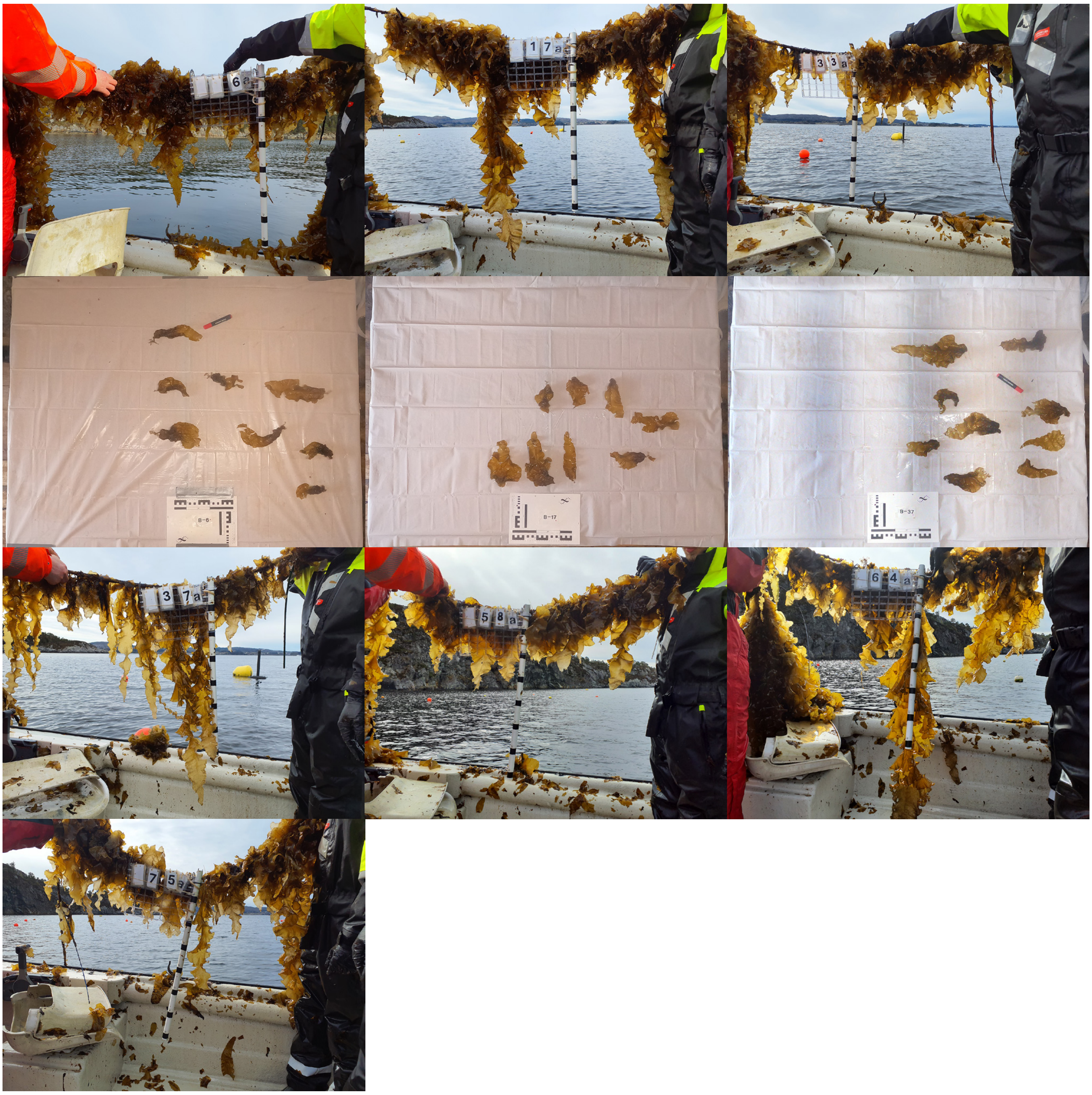

#### Genotype 9

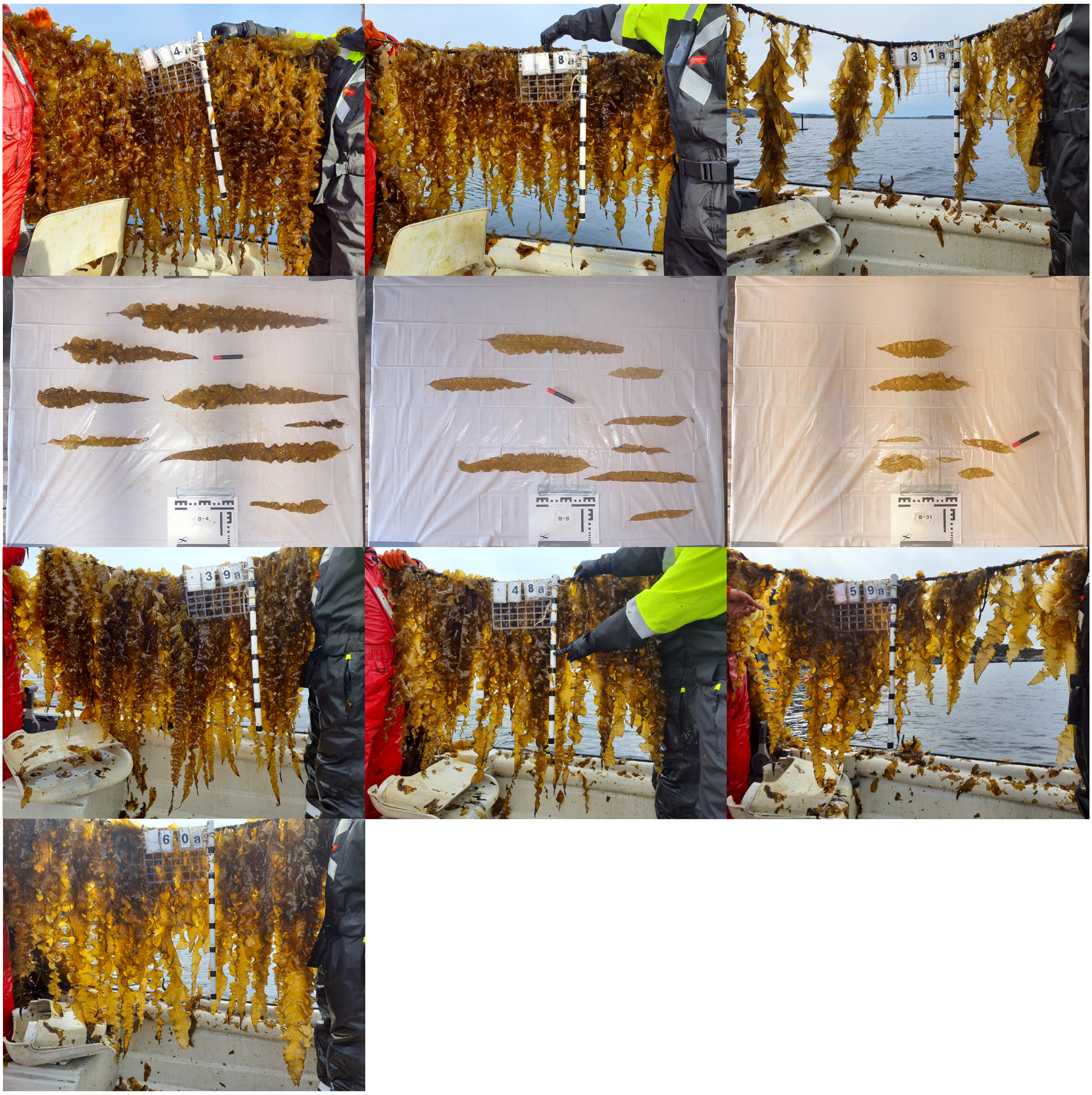

#### Genotype 11

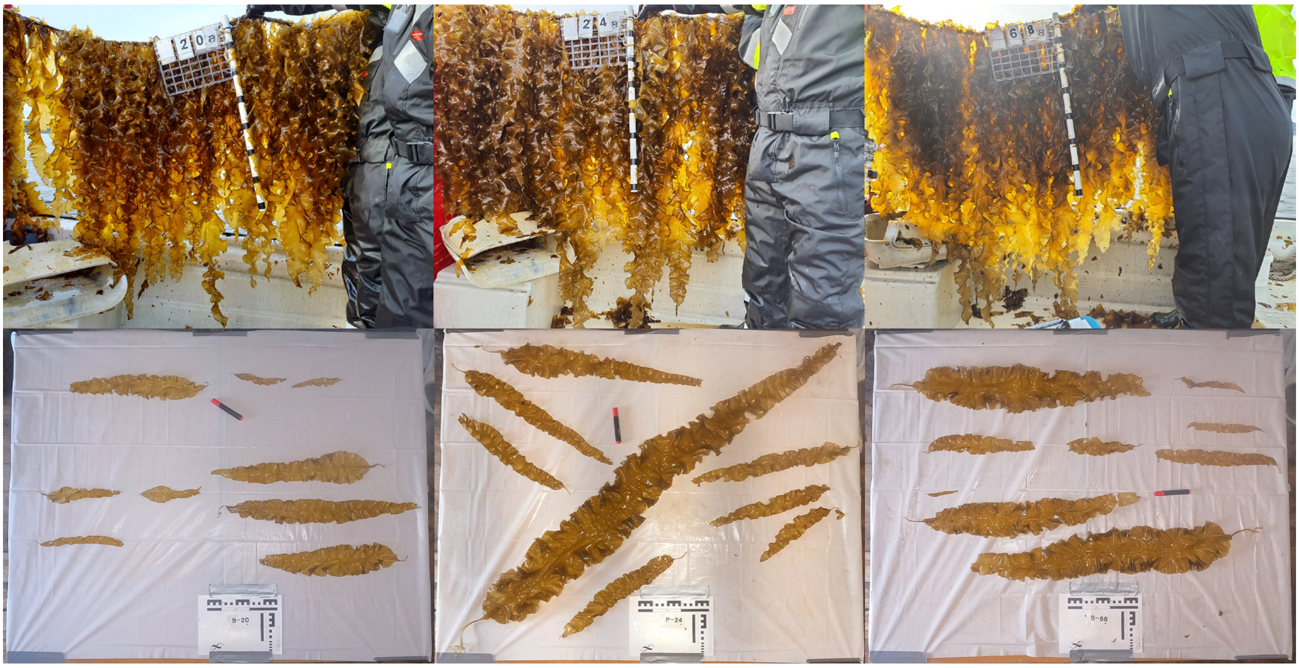

#### Genotype 11

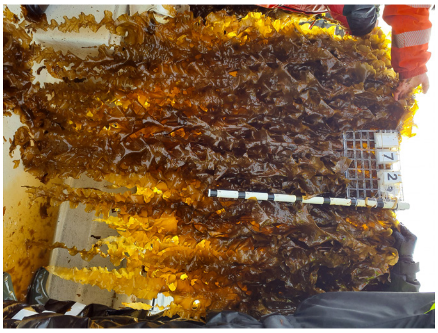

#### Genotype 12

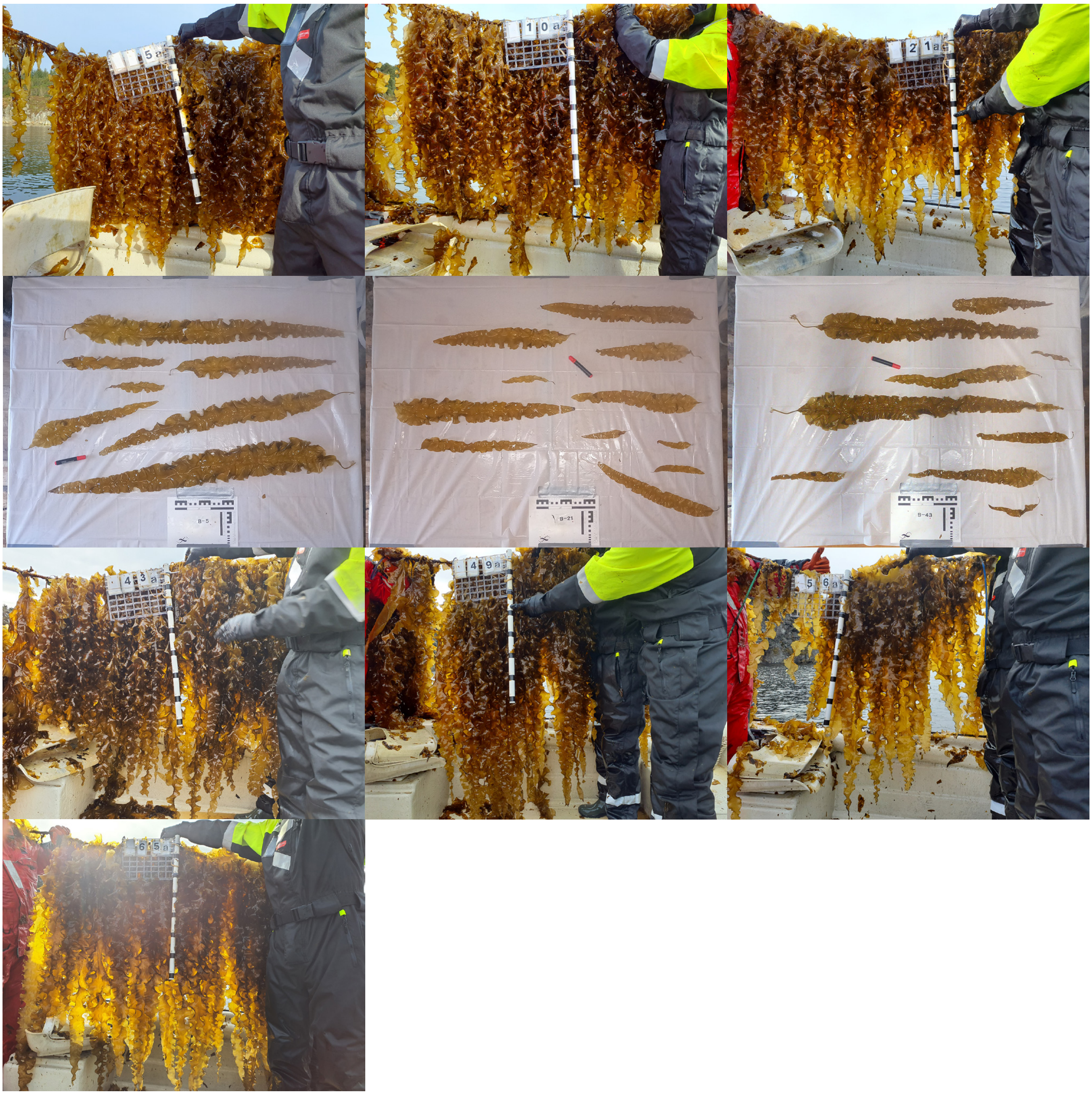
