## Supplementary figures and images for "Genotype and farm effects on yield and morphology reveal potential for breeding and site selection for sugar kelp"

### Figure S2

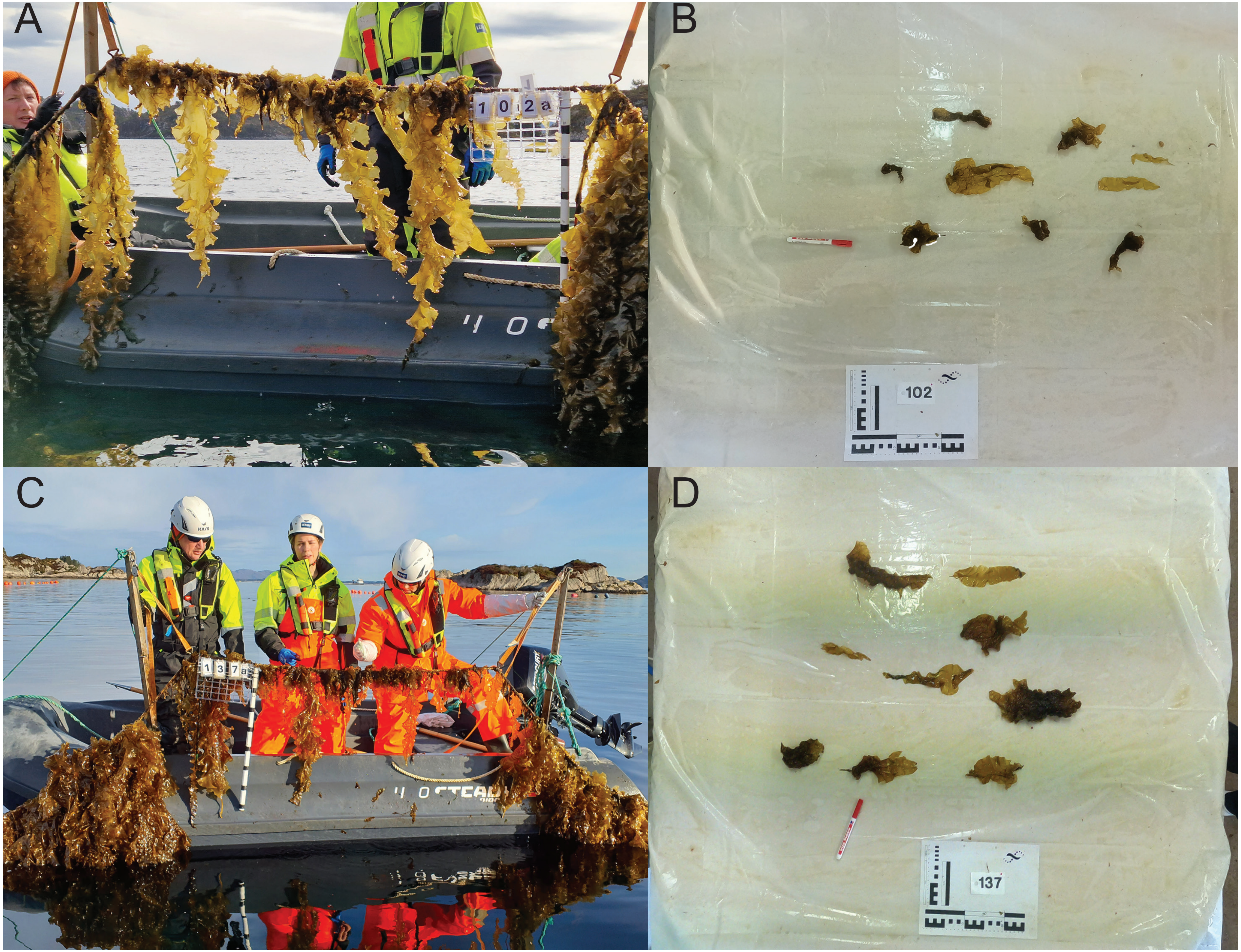

### Figure S3-1

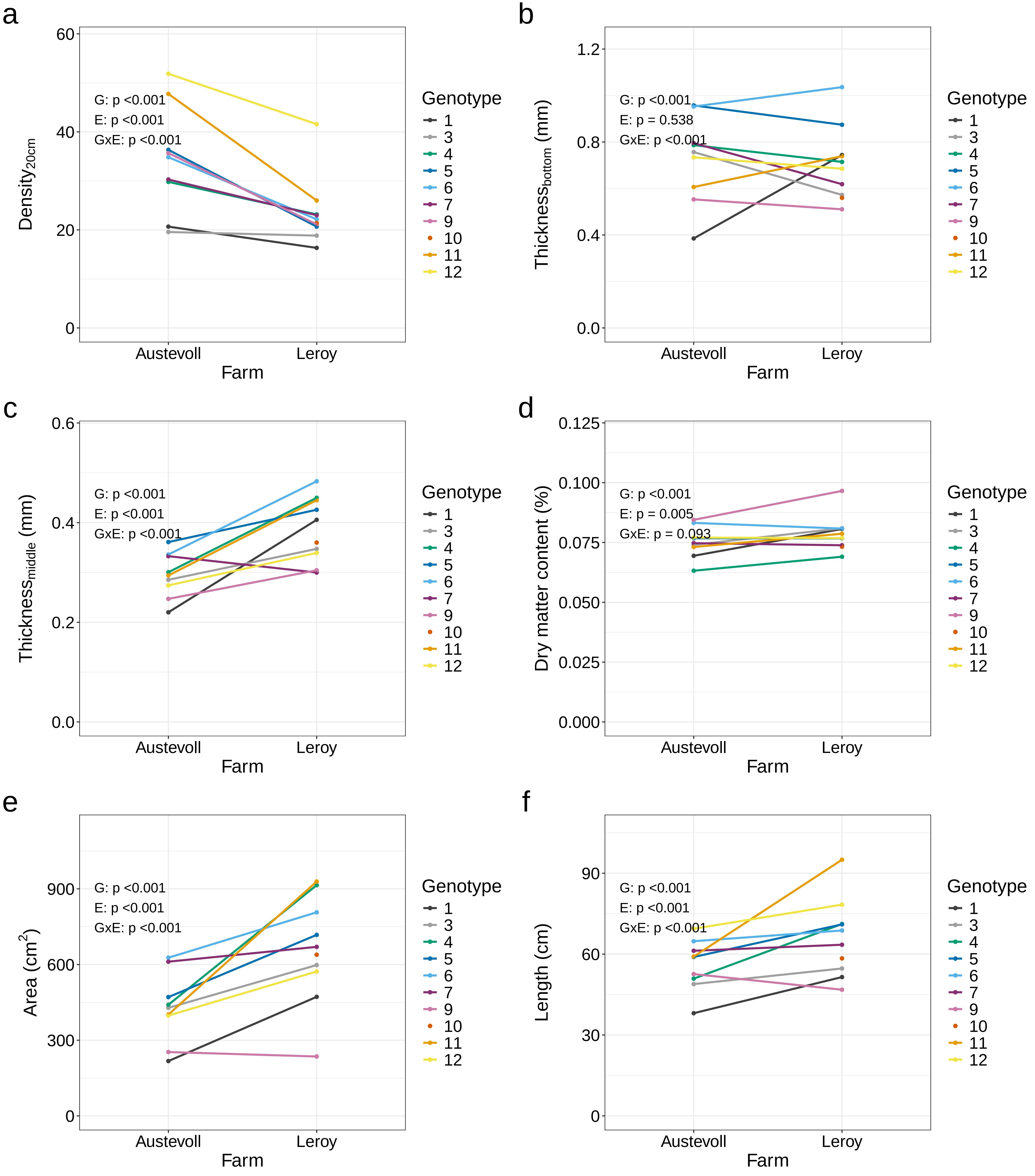

### Figure S3-2

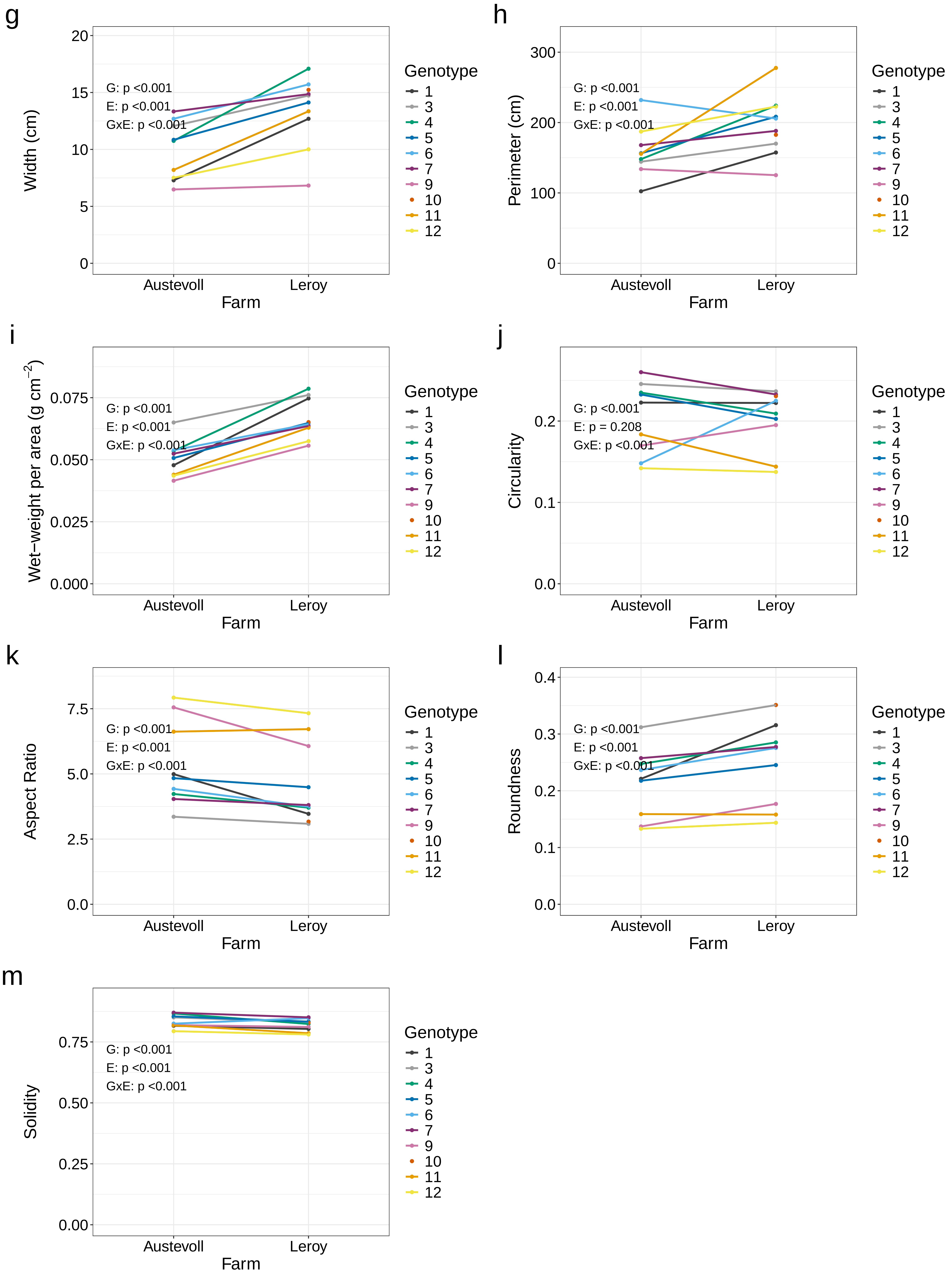

### Figure S4

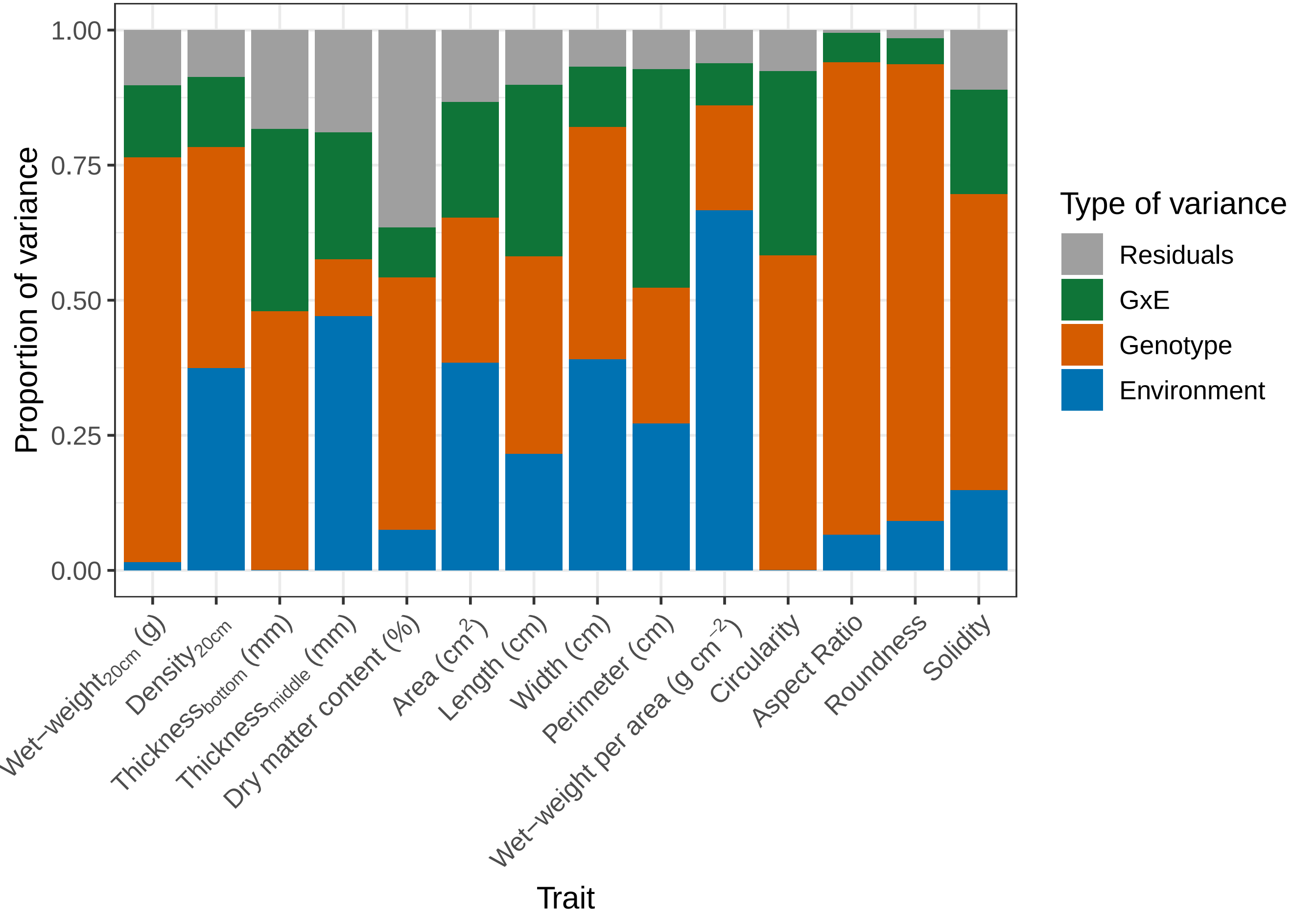

### Figure S5

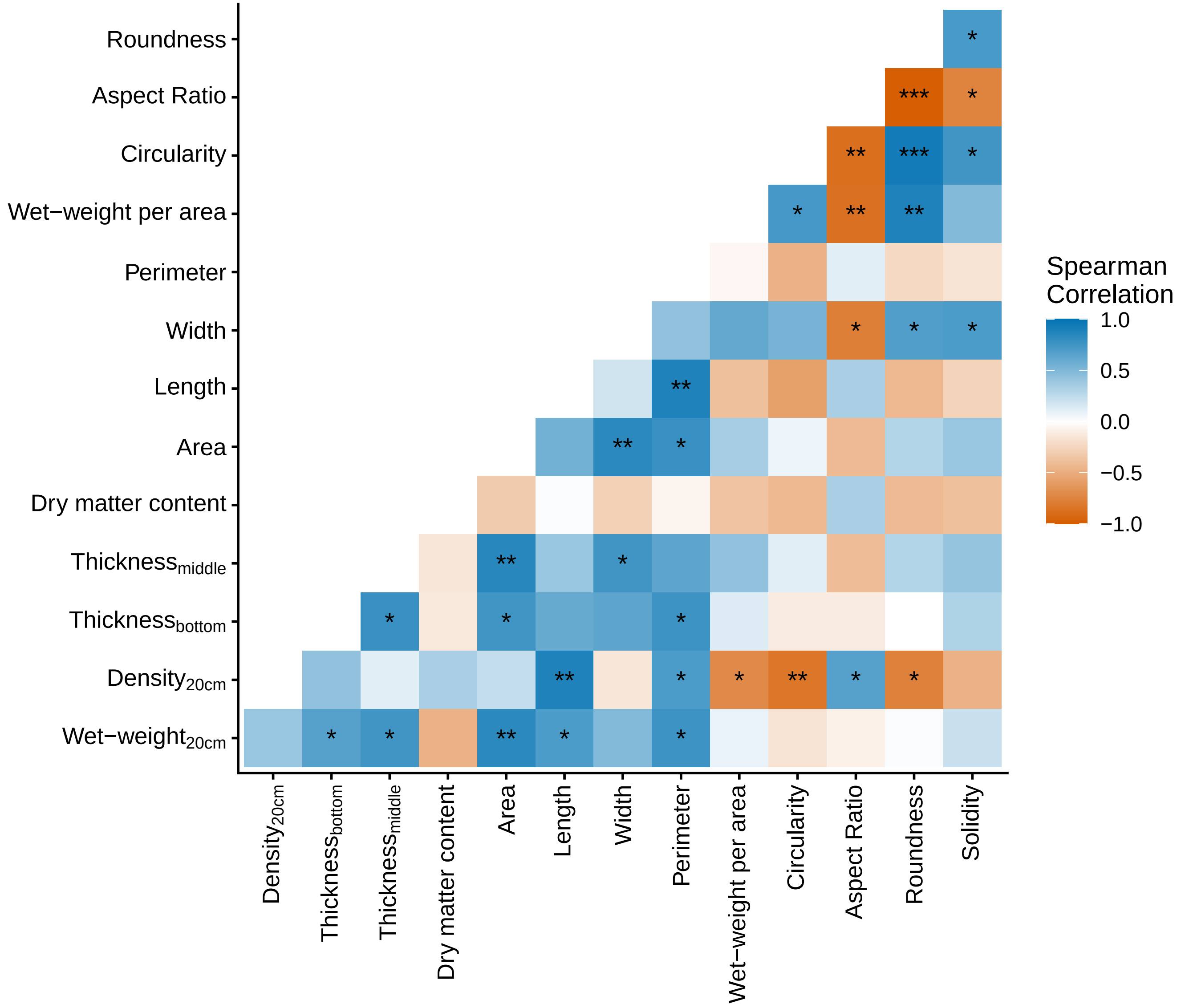
