## Supplementary material for "Genotype and farm effects on yield and morphology reveal potential for breeding and site selection for sugar kelp": Table S1

Table S2 Generalized heritability per farm.

|  | **Lerøy** | **Austevoll** |
| --- | --- | --- |
| **Trait** | **Heritability** | **Heritability** |
| Wet-weight_20cm_ (g) | 0.87 | 0.65 |
| Density_20cm_ | 0.80 | 0.83 |
| Thickness_bottom_ (mm) | 0.84 | 0.39 |
| Thickness_middle_ (mm) | 0.67 | 0.36 |
| Dry matter content (%) | 0.80 | 0.02 |
| Area (cm^2^) | 0.88 | 0.14 |
| Length (cm) | 0.90 | 0.00 |
| Width (cm) | 0.92 | 0.69 |
| Perimeter (cm) | 0.89 | 0.12 |
| Wet-weight per area (g/cm^2^) | 0.81 | 0.71 |
| Circularity | 0.92 | 0.40 |
| Aspect Ratio | 0.98 | 0.96 |
| Roundness | 0.97 | 0.95 |
| Solidity | 0.80 | 0.77 |
